# NanTex enables computational multiplexing and phenotyping of organelles across super-resolution modalities

**DOI:** 10.1101/2025.09.16.676574

**Authors:** Bela T.L. Vogler, Gregor J. Gentsch, Pablo Carravilla, Dominic A. Helmerich, Julia C. Heiby, Katharina Reglinski, William Durso, Teresa Klein, Felix Hildebrandt, Rohama Zahid, Katrin Spengler, Regine Heller, Christoph Kaether, Markus Sauer, Christian Eggeling, Christian Franke

## Abstract

Super-resolution microscopy (SRM) enables nanoscale visualization of cellular organelles but remains constrained by spectral overlap, labeling requirements, and temporal offsets in multicolor imaging. We present *NanTex*, a deep learning framework that introduces nanotexture as a universal descriptor of subcellular organization and achieves probabilistic demixing of multiple organelles from single-channel SRM images. Unlike segmentation methods that enforce exclusivity, *NanTex* preserves overlapping morphologies, enabling faithful reconstruction even in crowded regions. Trained on small curated datasets with augmentation, *NanTex* generalized across SMLM, MINFLUX, STED, SIM, and live-cell Airyscan, with modality-specific retraining where necessary. Demonstrated on cytoskeletal, endomembrane, and metabolic organelles, *NanTex* achieved high-fidelity reconstructions and extended uniquely to live-cell imaging, where it eliminated temporal misalignment and enabled quantitative tracking of vesicle-like ER subdomains. Notably, *NanTex* enables quantitative dynamic readouts directly from single-channel live-cell data, revealing and tracking hidden subdomains in real time without the need for additional labels. Beyond multiplexing, *NanTex* supported computational phenotyping by distinguishing structurally distinct yet molecularly identical populations, exemplified by nocodazole-induced microtubule depolymerization and its glyoxal-mediated modulation. Together, these results establish nanotexture as a paradigm-shifting descriptor for organelle identity and position *NanTex* as a modality-agnostic, label-efficient, and live-cell-ready strategy for quantitative nanoscale biology.

**Significance Statement:** *NanTex* reframes multiplexing as texture-based demixing, introducing nanotexture as a universal fingerprint of organelle identity. By enabling quantitative, label-efficient reconstructions across SRM modalities and live-cell phenotyping, including vesicle tracking from single-channel data, *NanTex* opens a path to studying organelle remodeling and disease progression with unprecedented fidelity across SRM modalities.

## Introduction

Understanding how organelles organize themselves and interact at the nanoscale is central to cell biology. However, the diffraction limit long obscured these structures. Fluorescence-based super resolution microscopy (SRM) has transformed biological imaging by enabling the visualization and quantitative analysis of subcellular structures at nanometer resolution^1,2^. Techniques such as single-molecule localization microscopy (SMLM)^3–6,15^, stimulated emission depletion (STED) microscopy^7^, MINFLUX microscopy^8^, structured illumination microscopy (SIM)^9,10^, and Airyscan microscopy^11^ have become indispensable tools for interrogating organelle architecture, dynamics, and interactions. Despite these advances, multiorganelle imaging remains fundamentally limited by the physics of fluorophores and acquisition. Spectral overlap, finite dye palettes, and the photophysical requirements of SRM^12,44^ restrict simultaneous labeling, while live-cell imaging faces additional constraints from phototoxicity and temporal misalignment^2^.

To address these bottlenecks, several multiplexing strategies have been proposed. Spectroscopic approaches such as fluorescence lifetime imaging microscopy (FLIM)^13^ and polarisation-resolved imaging^14^ extend color channels but remain restricted to a handful of species and are challenging to integrate across modalities. DNA-PAINT and Exchange-PAINT leverage iterative binding of labels to achieve large multiplexing capacity^15,16^, yet are inherently sequential and time-consuming, limiting their applicability to both dynamic and high-throughput imaging.

Complementary computational segmentation methods including *Cellpose*^17,18^, *Segment Anything*^19^, and large-scale learning frameworks^20^ all assume that structures can be discretely separated. In crowded regions where organelles overlap extensively these methods fail to preserve the overall nanoscale composition critical for biological interpretation.

Here, we introduce *NanTex* (from “*nanotexture*”), a content-agnostic computational multiplexing framework that leverages nanoscale textural features as a generalizable structural fingerprint for organelles. Rather than segmenting discrete objects, *NanTex* performs probabilistic demixing through deep learning and thereby enabling the reconstruction of multiple identically labeled organelles from single-channel, monochromatic SRM images. This paradigm shift reframes multiplexing as a problem of organelle-specific demixing rather than spectral separation or structural delineation. Importantly, *NanTex* is data-efficient, requiring ∼10 images per organelle for training when combined with augmentation^26,37^, and broadly generalizes across SRM modalities, including SMLM, MINFLUX, STED, SIM, and live-cell Airyscan.

We validate *NanTex* across fixed-cell and live-cell contexts, demonstrating accurate demixing of combinations of identically labeled microtubules, actin, clathrin, ER, mitochondria, peroxisomes, endosomes, lysosomes and vesicle-like ER-substructures even in crowded environments and challenging labeling conditions. Importantly, *NanTex* achieves live-cell monochromatic multiplexing, eliminating spectral cross-talk and temporal offsets that limit conventional multicolor imaging. For example, we applied *NanTex* to live-cell Airyscan data of ER–lysosome dynamics, building on our recent findings that implicate lysosome–ER contact sites of novel identified lysosomal proteins^21^.

Beyond multiplexing, *NanTex* uniquely supports computational phenotyping by disentangling structurally distinct but molecularly identical populations, exemplified by microtubule depolymerization and its glyoxal-mediated delay^33,34^. Together, our results establish *nanotexture* as a universal descriptor of organelle identity and *NanTex* as a paradigm-shifting tool for label-efficient and live-compatible structural biology that could enable quantitative nanomorphological analysis of cellular states, treatments, or disease perturbations.

## Results

### Haralick feature extraction identifies nanoscale organelle-specific textures

Cellular organelles exhibit various morphologies across the micrometer to molecular scales, accessible by SRM^23^. We reasoned that sufficiently labelled and resolved organelles will possess an intrinsic, characteristic and self-similar nanometric fingerprint, i.e. a *nanotexture*. To demonstrate the extraction of quantifiable nanoscale textures from SRM images that could enable the differentiation of multiple organelles, we extracted classical Haralick textural descriptors^24^ from single-molecule localization microscopy (SMLM) images of microtubules and endosomes, labelled by nanoparticles^47,48^ (**see methods, Fig. 1**). Using 26 features (mean and range values of 13 Haralick descriptors) computed in a sliding 7×7 pixel window, we observed partially separable distributions, with contrast-dependent features providing the clearest distinction (**Fig. 1b–c**). However, single-feature thresholding failed in overlapping regions, where textures intermix at the pixel scale (**Fig. 1d**). While this generally demonstrates the validity of an organelle-specific nanotexture, this classical segmentation approach failed to account for overlapping structures with different identities, inherent to most fluorescence imaging methods.

**Figure 1.**
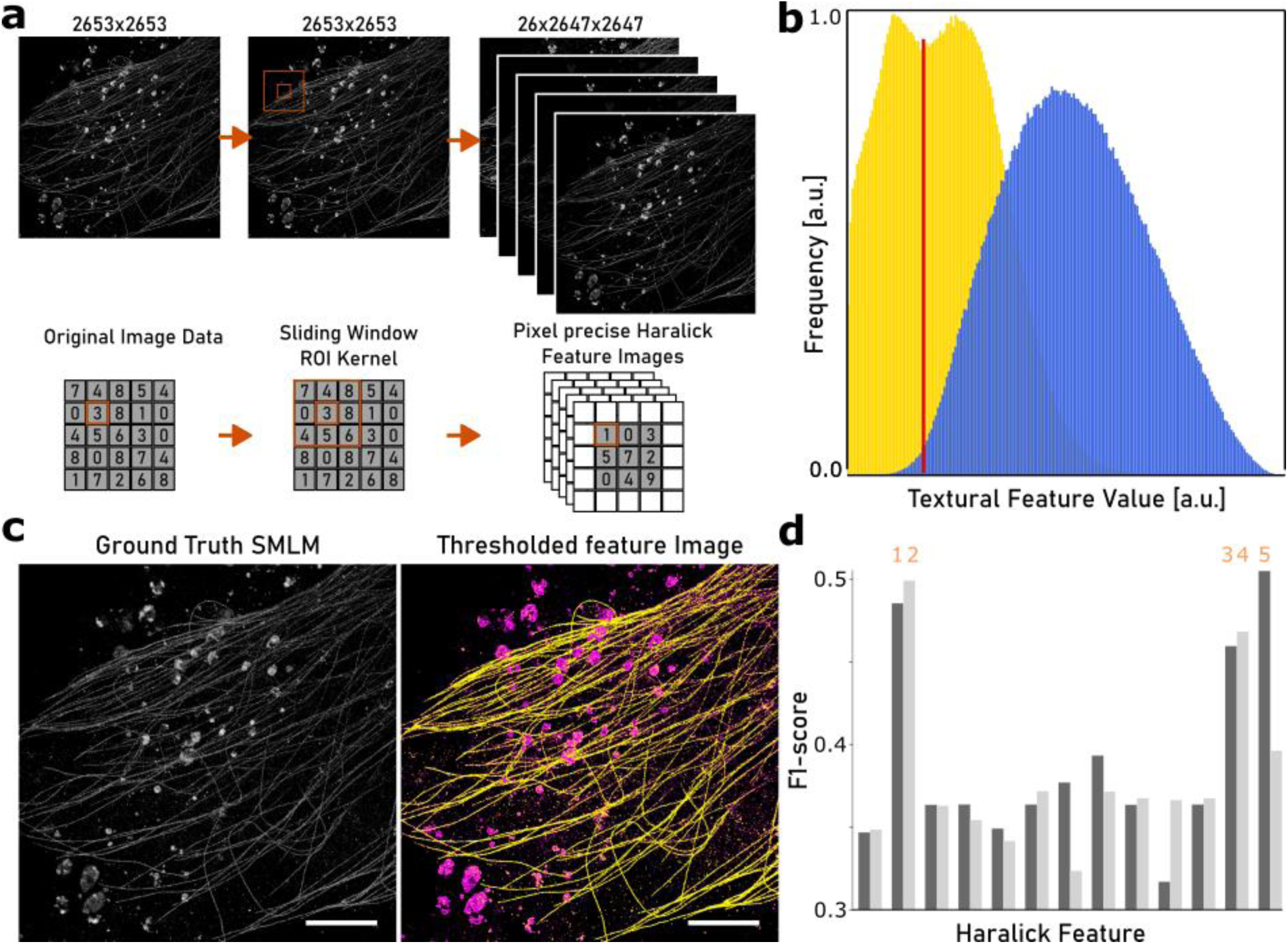
| Haralick feature extraction identifies nanoscale organelle-specific textures. (**a**) Workflow for pixel-precise Haralick feature extraction. Single-molecule localization microscopy (SMLM) images (here, microtubules and endosomes) were subdivided into sliding-window regions of interest (ROIs), and 26 Haralick texture descriptors were computed for each ROI, yielding feature images encoding nanoscale structural information. (**b**) Example distribution of single Haralick descriptors, showing partial separation in mean contrast (yellow) and continuous distribution in mean correlation (blue). A simple threshold (red line) allows only limited discrimination. (**c**) Representative ground truth SMLM image (left) and the corresponding thresholded Haralick feature image (right), where endosomes (magenta) and microtubules (yellow) are partially separated but still exhibit misclassification in overlapping regions. (**d**) Quantitative performance of all 26 Haralick features expressed as F1-scores for organelle classification. Only a subset of descriptors exceeded an F1-score of 0.4, namely Angular Second Moment (F1 ≈ 0.50, marked *1*), Contrast (F1 ≈ 0.48, marked *2*), Correlation (F1 ≈ 0.41, marked *3*), Sum of Squares Variance (F1 ≈ 0.45, marked *4*), and Homogeneity (F1 ≈ 0.52, marked *5*). These features showed moderate discriminative power, while the majority of descriptors performed substantially lower, highlighting that threshold-based approaches are insufficient for robust multiplexing. Scale bars: 5 µm.

### NanTex enables robust probabilistic demixing of monochromatic multi-organelle in silico SMLM data

Addressing the limitations of classical segmentation approaches, we developed *NanTex*, a machine-learning-enabled computational multiplexing framework leveraging the previously described organelle-specific nanoscale textures. Unlike segmentation methods, *NanTex* utilizes probabilistic demixing enabled by a U-Net^25^ trained to reconstruct multiple organelle output channels from monochromatic SRM input images (**see methods**). In short, the model employs a standard encoder-decoder architecture, with each convolutional block followed by batch normalization and ReLU activation. A 1×1 convolutional layer using linear regression outputs separate probabilistic predictions for each organelle. It is trained to produce the most probable component images contained within a superimposed multi-organelle grayscale input. Importantly, this generative approach preserves intensity and nanoscale nuances of overlapping organelles, fundamentally distinguishing *NanTex* from segmentation-based multiplexing methods. Being able to leverage heavy data augmentation (**see methods**)^26^, the network can be efficiently trained with ∼10 images per organelle and SRM modality, a requirement easily fulfilled by most SRM modalities and users.

We first demonstrated *NanTex*’s multiplexing capability using SMLM datasets with highest structural resolution, comprising three organelles often used in SRM benchmarking with distinct but overlapping structural profiles: microtubules, actin, and endosomes (i.e. vesicles)^27,28^. *NanTex* was trained on a stack of computationally superimposed and augmented sets of organelles (sourced partially from shareloc^42^, **see methods**) with the task of re-generating the individual underlying images. This allowed for the successful multiplexing of different, *in silico* combined, multi-structural datasets and benchmarking of the network.

Globally, *NanTex* reconstructed all three overlapping individual structures of the test set, i.e. microtubules, endosomes, and actin, with high fidelity, quantified by structural (multi-scale) self-similarity index measure^29–31^ (SSIM = 0.80 - 0.87; MS-SSIM = 0.90 - 0.92, **Fig. 3, see Table 1**). Importantly, multiplexing performance even rose within densely overlapping ROIs (SSIM = 0.84 - 0.90; MS-SSIM = 0.93 - 0.94, **Fig. 3**), where conventional segmentation completely failed to separate signals. While SSIM and MS-SSIM are full-reference, perception-informed metrics that assess similarity in luminance, contrast, and structure, and are therefore designed to correlate with human judgments of image quality (SSIM ranges from -1 to 1, with 1 indicating identical images), MS-SSIM in particular captures image fidelity across multiple spatial scales, providing greater alignment with perceptual quality than single-scale SSIM (**see Table 1, Supplementary Discussion, Supplementary** Fig. 1). Accordingly, the consistently high MS-SSIM values observed here (0.90–0.94) indicate near-perfect, perceptually indistinguishable reconstruction of the ground truth. These results highlight *NanTex*’s unique ability to demix at least three spectrally identical and structurally overlapping organelles from a single monochromatic image, fundamentally surpassing segmentation-based approaches.

**Figure 2.**
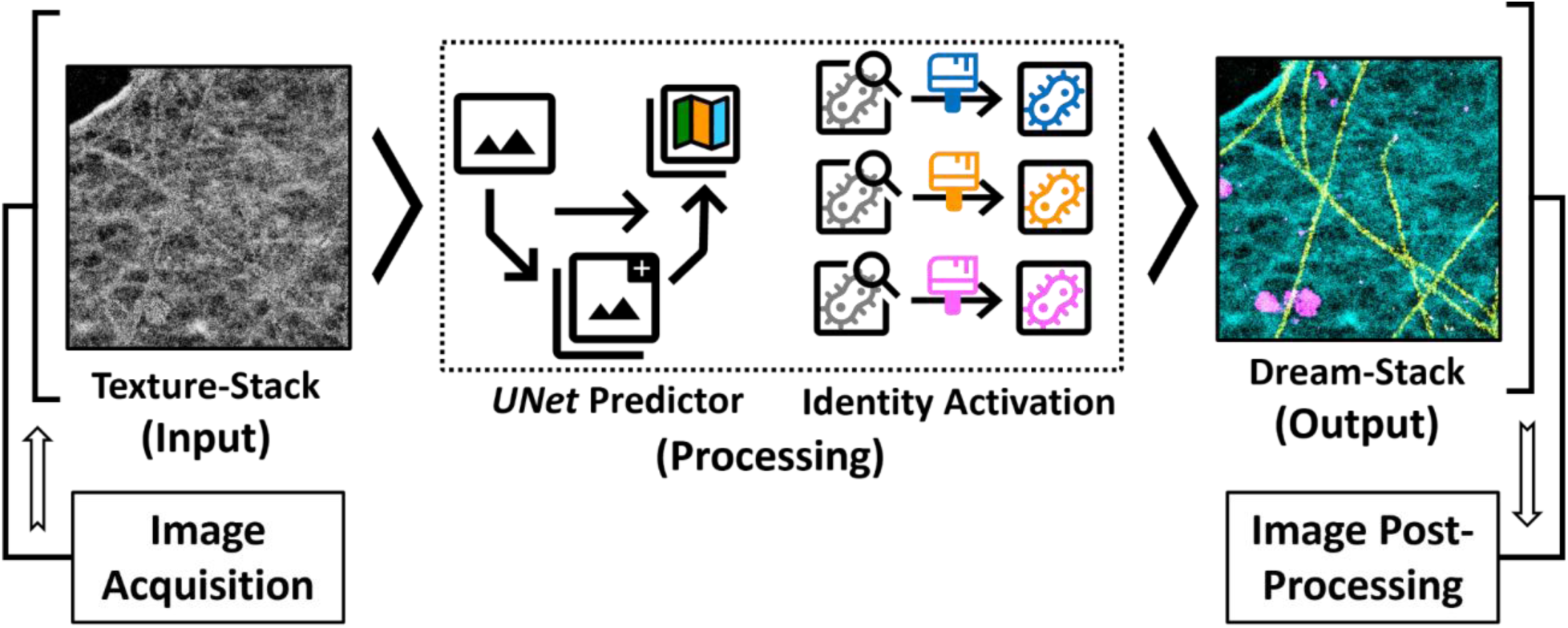
| *NanTex* architecture and probabilistic demixing workflow. Schematic overview of the *NanTex* pipeline. Monochromatic SRM acquisitions (left) containing multiple organelles are subdivided into training patches and passed through a U-Net–style encoder–decoder network (middle). The network employs probabilistic demixing, directly reconstructing independent organelle channels rather than performing classical segmentation. Color-coded icons illustrate augmentation and network operations applied during training (rotation, flips, Gaussian noise/blur, gamma correction), which ensure robustness to variability in labeling density, photophysics, and imaging artifacts. The output (right) is a set of multiplexed organelle-specific channels that preserve both intensity and fine structural detail, here shown as ER (cyan), microtubules (yellow), and clathrin (magenta). This architecture enables accurate demixing even under high structural overlap, with training requirements of ∼10 input images per organelle and modality, augmented into several thousand training patches.

**Figure 3.**
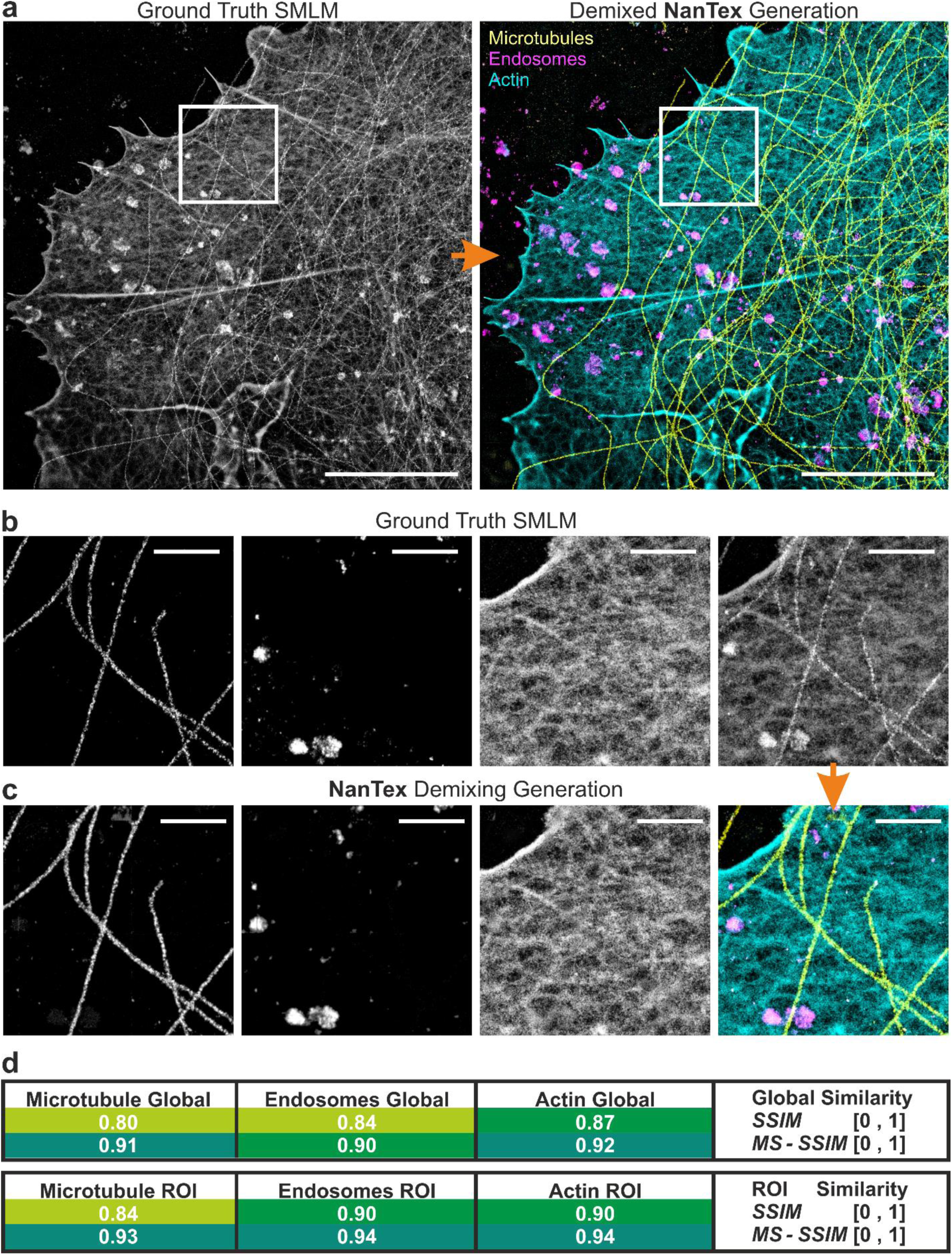
| *NanTex* enables high-fidelity organelle specific multiplexing of triple-identity monochromatic SMLM data in silico. (**a**) Exemplary SRM image rendered from unrelated SMLM data of microtubules, actin and endosomes, combined in-silico in native greyscale (*left*) and its demixed multi-colour counterpart after *NanTex* analysis in false colours (*right*, yellow: microtubules, cyan: actin, magenta: endosomes). (**b**) Magnified views of the region indicated in (**a**), exhibiting significant content in all three ground-truth channels (*from left to right:* microtubules, endosomes, actin, sum), in comparison with (**c**) their respective counterparts after *NanTex* analysis. (**d**) Quantitative evaluation of *NanTex* reconstructions. Structural similarity (SSIM) and multi-scale SSIM (MS-SSIM) were computed globally across full fields of view (**a**) and locally in regions of dense overlap (**b-c**). *NanTex* achieved high global fidelity for all three organelles (microtubules: SSIM = 0.80, MS-SSIM = 0.91; endosomes: SSIM = 0.84, MS-SSIM = 0.90; actin: SSIM = 0.87, MS-SSIM = 0.92) and further improved similarity within overlapping ROIs (SSIM = 0.84–0.90, MS-SSIM = 0.93–0.94). These consistently high MS-SSIM values indicate near-perceptual indistinguishability from the ground truth, in stark contrast to the failure of intensity thresholding to recover overlapping structures. Scale bars: 5 µm (a), 1 µm (b-c).

**Table 1.** | Guidelines for interpretation of SSIM and MS-SSIM scores in *NanTex* benchmarking

| SSIM | MS-SSIM | Interpretation | Practical meaning for SRM demixing |
| --- | --- | --- | --- |
| $\geq 0.95$ | $\geq 0.95$ | Virtually indistinguishable from ground truth | Nanotexture and morphology fully preserved; reconstructions visually identical. |
| 0.90–0.95 | $\geq 0.90$ | Excellent fidelity | Minor smoothing, but organelle assignment and morphology reliable for quantitative analysis. |
| 0.80–0.90 | $\geq 0.85$ | Good fidelity | Structures correctly demixed; local deviations acceptable. Suitable for comparative studies. |
| 0.70–0.80 | $\geq 0.80$ | Moderate fidelity | Blur or local misassignments visible, but overall shape preserved. Use with caution for quantification. |
| $< 0.65$ | $< 0.70$ | Poor fidelity | Major loss of texture or wrong assignment; reconstruction unsuitable for biological interpretation. |

Encouraged by these in silico results, we next asked whether *NanTex* could achieve comparable multiplexing fidelity on experimental *d*STORM acquisitions of identically labeled organelles.

### Experimental validation of monochromatic SMLM multiplexing with NanTex

To validate *NanTex* in realistic contexts, we acquired monochromatic *d*STORM datasets in which distinct organelles were deliberately labeled with the same fluorophore (AF647), resulting in composite images that appear structurally entangled. *NanTex* was then applied to demix these single-color overlays into separate channels corresponding to the original structures.

In the first experiment, we multiplexed three organelles (**Fig. 4**, yellow: microtubules, magenta: clathrin, cyan: ER) recorded as independent AF647 *d*STORM acquisitions to ensure ground truth quantifyability. *NanTex* faithfully separated the three structures, maintaining high fidelity over 500 test images (**see methods**), with mean SSIM/MS-SSIM values of 0.86/0.94 for microtubules, 0.83/0.86 for clathrin, and 0.83/0.89 for ER. A representative ROI (Fig. 4) shows slightly lower but still robust scores (microtubules: 0.82/0.94; clathrin: 0.67/0.77; ER: 0.75/0.85), consistent with the partially overlapping diffuse monomeric fractions of clathrin and ER, whereas the filamentous microtubules exhibited the highest fidelity. Together, these results confirm that *NanTex* preserves fine structural detail while tolerating biological complexity arising from shared molecular pools.

**Figure 4.**
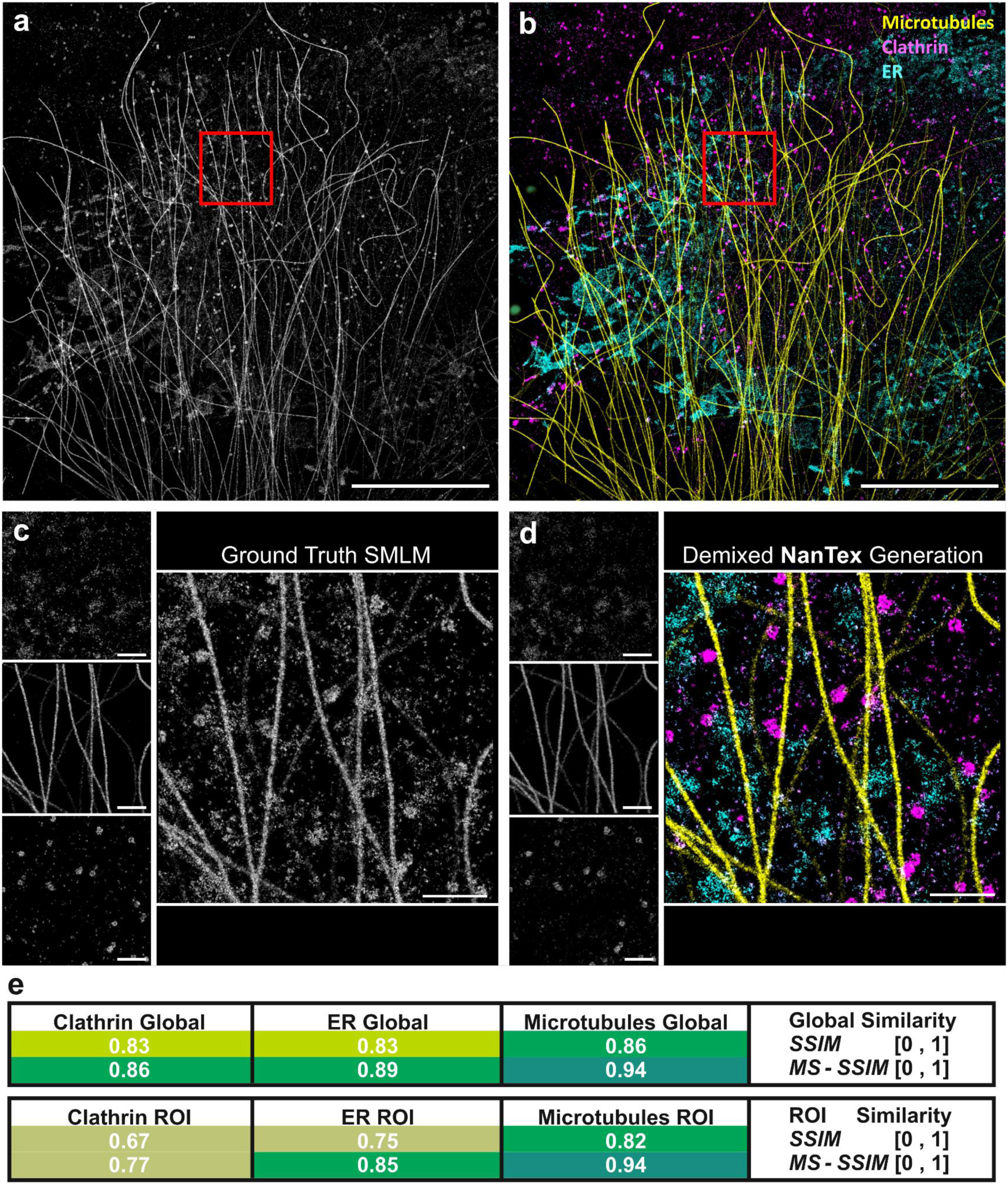
| *NanTex* demixes experimental SMLM overlays of identically labeled organelles. (**a**) Composite monochromatic *d*STORM image generated by combining individual AF647 acquisitions of microtubules, clathrin, and ER. (**b**) *NanTex* multiplexing results in false colours (yellow: microtubules, magenta: clathrin, cyan: ER). (**c**) Ground-truth channels of a randomly selected ROI. (**d**) Corresponding *NanTex* demixed channels and false-colour image of the same ROI. (**e**) Quantitative evaluation of the same ROI using SSIM and MS-SSIM, displayed in false colours. Global mean SSIM/MS-SSIM values were 0.86/0.94 for microtubules, 0.83/0.86 for clathrin, and 0.83/0.89 for ER; corresponding ROI values were 0.82/0.94, 0.67/0.77, and 0.75/0.85, respectively. The reduced scores for clathrin and ER reflect their partially overlapping diffuse monomeric fractions, while filamentous microtubules reached the highest fidelity. The representative example in (**a–e**), which was not cherry-picked, illustrates typical performance but yields slightly lower similarity scores than the overall dataset averages. Scale bars: 5 µm (a–b), 1 µm (c–d).

We next validated *NanTex* on pairwise monochromatic *d*STORM data (**Fig. 5**). For single-label microtubules and clathrin (AF647) that were imaged by *d*STORM simultaneously. *NanTex* successfully disentangled the cytoskeletal network from vesicular structures. Of note: clathrin-coated vesicles are consistently resolved despite spatial overlap with microtubules, without destroying their respective integrity. Importantly, the demixed microtubule channel shows virtually no unspecific background signal, consistent with the expected filamentous localization, while clathrin is robustly detected in both its vesicular and diffuse monomeric membrane fractions. For identically labeled microtubules and ER, *NanTex* also achieved a reliable separation, although the overall labeling efficiency for the combined structures was lower due to protocol mismatches. Despite this suboptimal staining, the remaining ER and microtubule network were accurately demixed, underlining the robustness of the model even in imperfect experimental conditions.

**Figure 5.**
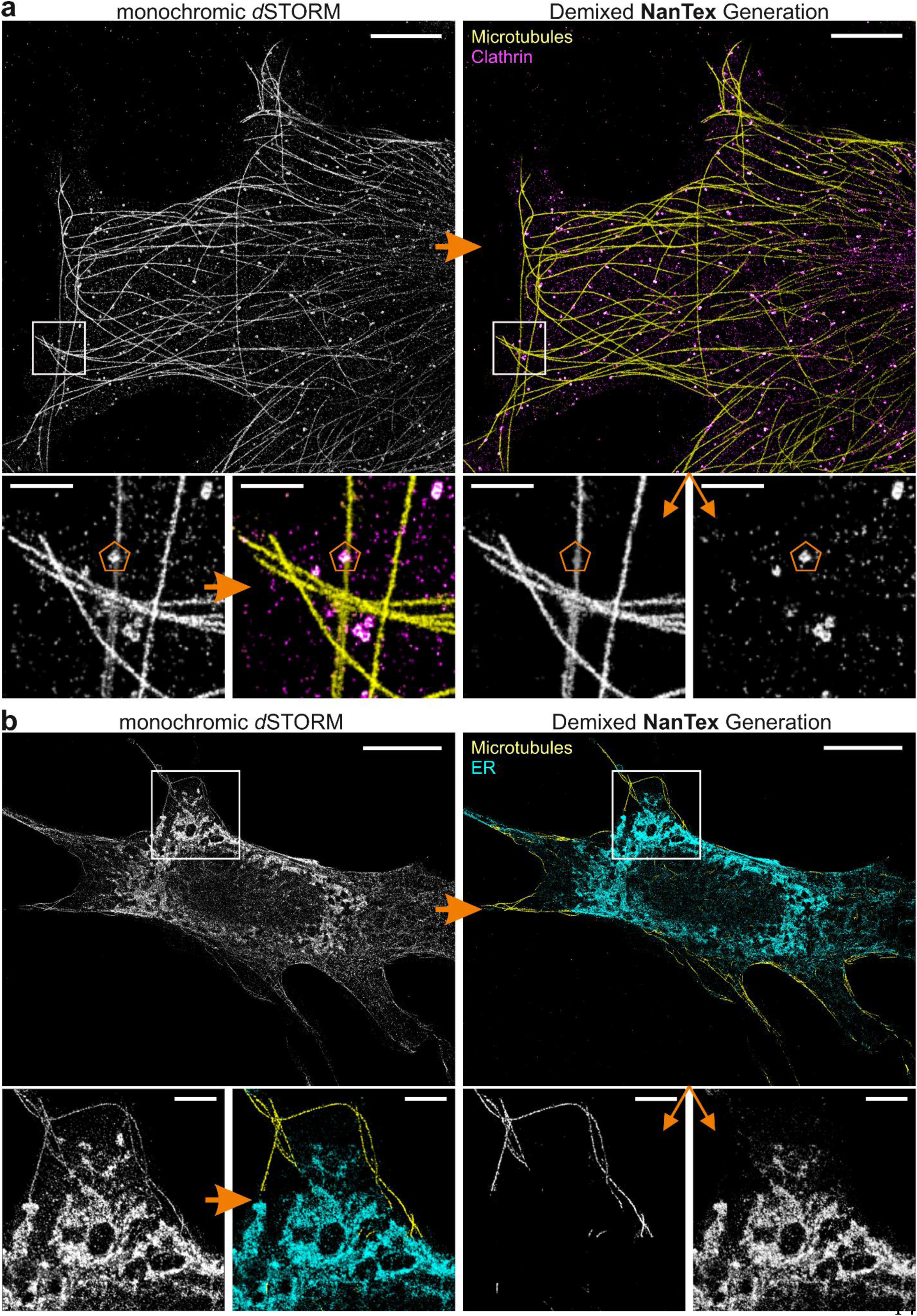
| Experimental validation of monochromatic SMLM multiplexing with *NanTex*. (**a**) Representative *d*STORM dataset of identically labeled microtubules and clathrin (AF647). Top row: experimentally acquired monochromatic input (*left*) and *NanTex* multiplexing result (*right*; yellow: microtubules, magenta: clathrin). Bottom row: zoom-in of the indicated ROI showing ground truth (left), *NanTex* false-color overlay, and the corresponding demixed channels for microtubules and clathrin. An orange hexagon highlights a region where a clathrin-coated vesicle is cleanly separated from an overlapping microtubule. (**b**) Equivalent *NanTex* analysis of identically labeled microtubules and ER (cyan). Top row: experimentally acquired monochromatic input and *NanTe*x multiplexing result. Bottom row: zoom-in ROI with ground truth, false-color overlay, and individual *NanTex* channels. While labeling efficiency for ER was suboptimal due to protocol mismatches, the remaining structures are still cleanly multiplexed from microtubules. Scale bars: 5 μm (overview), 1 μm (zoom-ins).

Together, these results establish that *NanTex* multiplexes monochromatic SMLM data across different organelles with high fidelity, preserves fine structural features, and tolerates experimental variability such as uneven labeling efficiency.

### NanTex demonstrates multiplexing capability across MINFLUX, STED, SIM, and Airyscan microscopy

Having validated *NanTex* for SMLM, we next tested its generalizability across other fundamental SRM modalities (**Fig. 6**), each with strengths and limitations: MINFLUX provides unmatched localization precision but is intrinsically low throughput^32^; conversely, SIM, STED, and Airyscan are widely accessible but operate at lower resolution, potentially constraining nanotexture detection.

**Figure 6.**
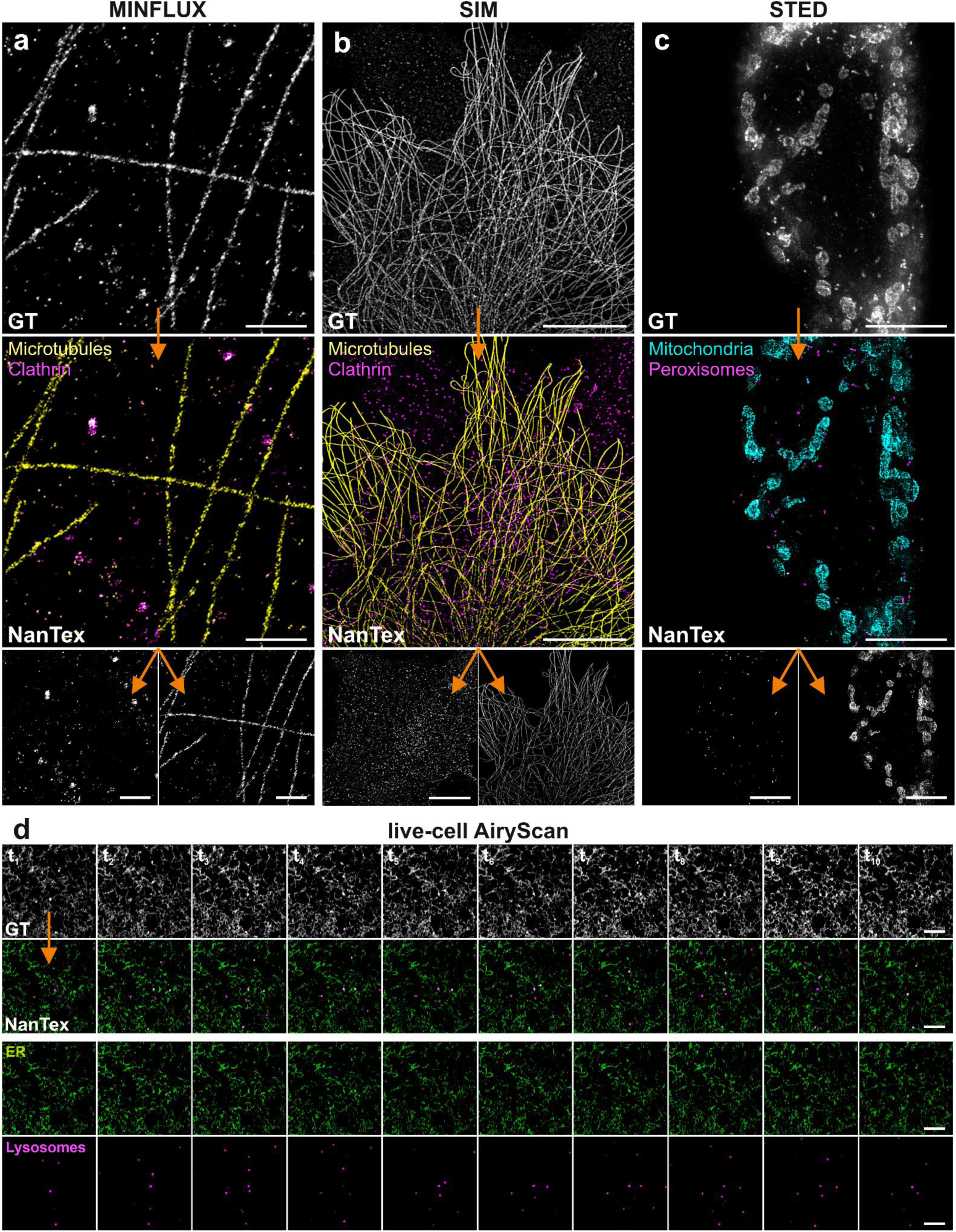
**| *NanTex* multiplexing capability across advanced SRM modalities**. (**a**) Representative MINFLUX images of identically labeled clathrin and microtubules (AF647). Top: ground truth grayscale acquisitions; middle: *NanTex* multiplexing result (yellow: microtubules, magenta: clathrin) obtained with an SMLM-trained model without retraining; bottom: individual demixed channels. (**b**) Representative SIM analysis of identically labeled microtubules and clathrin. Top: monochromatic input; middle: *NanTex* multiplexing; bottom: single demixed channels. *NanTex* SIM reconstructions achieved high global MS-SSIM values (0.91 for microtubules, 0.87 for clathrin). (**c**) STED multiplexing of mitochondria (Tom20, Abberior STAR Red, cyan) and peroxisomes (PMP70, Abberior STAR Orange, magenta). Top: monochromatic acquisitions; middle: *NanTex* multiplexing; bottom: demixed channels. Quantitative evaluation showed robust demixing with MS-SSIM = 0.86 (mitochondria) and 0.90 (peroxisomes), despite lower single-scale SSIM values (∼0.61–0.67), consistent with STED’s reduced SNR. (**d**) Live-cell Airyscan recordings of ER (GFP-ATL3, green) and lysosomes (LysoTracker, magenta). Top: ground truth grayscale frames from a representative time course; middle: *NanTex* multiplexing; bottom: demixed channels. *NanTex* reconstructions achieved near-perfect fidelity for lysosomes (SSIM = 0.96, MS-SSIM = 0.94), while ER reconstructions scored lower at single-scale (SSIM = 0.53) but higher across scales (MS-SSIM = 0.75), reflecting robust recovery of global ER morphology. The entire time course is shown in **Supplementary Video 1** and related ER-only Airyscan recordings (Supplementary Fig. 4, Supplementary Video 2) not only demonstrate vesicle demixing but also *NanTex*-enabled single-particle tracking of vesicular substructures, providing quantitative readouts of dynamic behavior. Scale bars: 1 μm (MINFLUX), 5 μm (SIM, STED, Airyscan).

Applying our SMLM-trained model directly to MINFLUX images of identically labeled clathrin and microtubules yielded accurate multiplexing (**Fig. 6a**), establishing a proof-of-concept for cross-modality generalization without retraining - a particularly promising result given the low data yield of MINFLUX microscopy. However, performance degraded when the same model was applied to blurred *d*STORM or SIM datasets (**Supplementary** Fig. 2), demonstrating that cross-modality *transfer* works when the modality’s effective resolution and/or contrast is close to the source domain.

We next trained *NanTex* directly on lattice SIM datasets of microtubules and clathrin. Despite the modest lateral resolution (∼120 nm at 640 nm excitation), reconstructions achieved high MS-SSIM (0.91 for microtubules, 0.87 for clathrin), demonstrating that organelle-specific nanotextures can be recovered even under resolution-limited conditions (**Fig. 6b**). For STED, similar samples yielded poor performance (**Supplementary** Fig. 3), consistent with the subpar STED utility of AF647. However, optimized dual-color STED imaging of mitochondria (Abberior STAR Red) and peroxisomes (Abberior STAR Orange) enabled robust multiplexing (MS-SSIM = 0.86 and 0.90; **Fig. 6c**), despite modest single-scale SSIM values (∼0.61–0.67). This demonstrates that *NanTex* faithfully preserves global organelle structure under conditions of sufficient dye brightness and contrast, and further underscores the utility of multi-scale quality metrics in lower-SNR modalities.

#### NanTex enables multiplexed live-cell imaging

Encouraged by the strong SIM results, we next focused on live-cell applications. Compared to SIM, Airyscan microscopy offers similar resolution, but superior temporal performance and robustness making it the workhorse for dynamic multicolor recordings. Using U2OS cells expressing GFP-ATL3 and stained with LysoTracker, *NanTex* cleanly demixed ER and lysosomes from single-channel overlays (**Fig. 6d**, **Supplementary Video 1**). Quantitatively, fidelity was near-perfect for lysosomes (SSIM = 0.96, MS-SSIM = 0.94) and robust for ER (SSIM = 0.53, MS-SSIM = 0.75), reflecting strong recovery of global morphology despite lower pixel-level contrast.

Our data directly connects to our recent findings on newly discovered lysosomal proteins^21^ and the emerging importance of ER–lysosome interactions in cellular physiology and pathology. In particular, ER–lysosome contact sites are increasingly recognized as key regulators in a wide range of disorders, where subtle perturbations of organelle communication underlie disease phenotypes.

Importantly, *NanTex* also revealed previously hidden heterogeneity within single-labeled datasets. In Airyscan recordings of ER-only mRFP-KDEL U2OS cells, vesicle-like substructures became apparent that could not be captured by segmentation or orthogonal labeling, but were cleanly separated from the reticular ER by *NanTex*, consistent with ER-exit sites or sites of ER-phagy (**Supplementary** Fig. 4, **Supplementary Video 2**). In addition to qualitative separation, *NanTex* enabled quantitative analysis of the demixed vesicle channel. Using *TrackMate*-based single-particle tracking^49^, we obtained mean-squared displacement (MSD) curves of vesicle-like structures across time-lapse recordings (**see methods**, **Supplementary** Fig. 4e)^50^. The resulting diffusion coefficients extracted from MSD (time-average: 0.013±0.07 μm²/s, ensemble-average: 0.013±0.01 μm²/s) indicated a slow and homogeneous (ergodicity: 1.0) diffusion, while the anomalous diffusion alpha coefficients (time-average: 0.76±0.24, ensemble-average: 0.85) suggested confined movement, consistent with ER-bound compartments. This demonstrates that *NanTex* not only unmasks hidden structural subdomains but also renders them accessible to quantitative dynamic readouts directly from monochromatic live-cell data. This ability to uncover and track structurally distinct subdomains despite their identical label in real time, establishes live-cell imaging as one of the strongest potential application areas for *NanTex*. This could enable direct computational phenotyping of organelle heterogeneity and remodeling upon disease and treatment in real time in future studies.

### NanTex enables computational phenotyping of structurally distinct molecular populations

Building on the cross-modality results, we next asked whether *NanTex* could be extended beyond separating different organelles to disentangling structural heterogeneity within a single molecular population. If successful, *NanTex* could also serve as a tool for *computational phenotyping*, i.e. detecting and quantifying structurally distinct subpopulations within otherwise indistinguishable molecular pools, inaccessible by orthogonal labelling strategies.

To establish that *NanTex* recognizes structural nanotexture rather than spurious intensity cues, we conducted a controlled downsampling experiment on *d*STORM microtubule datasets, rendered with successively fewer localizations. Tubulin remained molecularly present but the filamentous nanotexture degraded into diffuse signal, leading to a sharp breakdown of SSIM and MS-SSIM values (**Supplementary** Fig. 5), validating that *NanTex* faithfully recognizes the self-similar tubular structure itself, not auxiliary metadata.

Guided by these controls, we applied *NanTex* to microtubule depolymerization phenotypes in human umbilical vein endothelial cells (HUVECs) treated with nocodazole^33^, with or without glyoxal supplementation at non-fixative concentrations^22,34^. Using the SMLM-trained model (microtubules, ER, clathrin), *NanTex* demixed intact filamentous microtubules (yellow) from diffuse tubulin pools (magenta) despite their identical antibody labeling (**Fig. 7 a-h**). The resulting per-cell polymerization index (PolIndex, **Fig. 7 i, see methods**) distributions revealed significant and quantifiable phenotypes, i.e. drastic loss of filament integrity under nocodazole treatment (PolIndex = −0.61 ± 0.25) and a significant trend toward delayed depolymerization in the presence of both, nocodazole and glyoxal (PolIndex = 0.16 ± 0.23), while glyoxal alone resulted in a modest reduction (PolIndex = 0.42 ± 0.30) relative to untreated cells (PolIndex = 0.65 ± 0.22).

**Figure 7.**
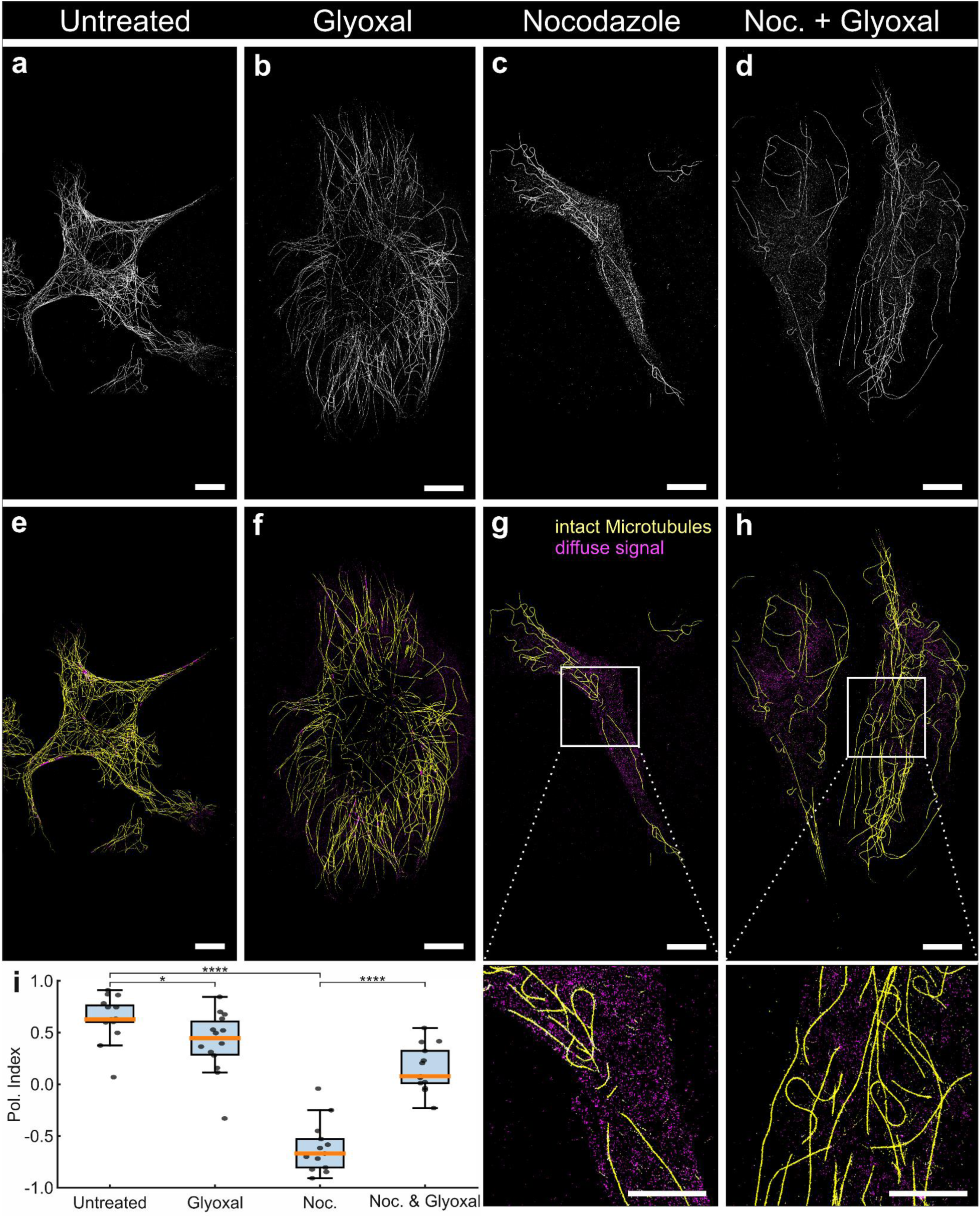
| *NanTex* enables computational phenotyping of microtubule depolymerization in HUVECs. (**a–d**) Representative *d*STORM images of α-tubulin–labeled microtubules (AF647) in primary HUVECs under four conditions: untreated, glyoxal-stabilized (1 mM, 48 h), nocodazole-treated (10 µM, 5 min), and combined nocodazole + glyoxal treatment. (**e–h**) Corresponding *NanTex* multiplexing results, separating intact filamentous microtubules (yellow) from diffuse depolymerized tubulin (magenta). Insets (**g,h**) highlight representative regions where *NanTex* resolved diffuse signals and filaments despite their identical label. (**i**) Boxplots with jittered single-cell points of the polymerization index (PolIndex) for Untreated (n = 15), Glyoxal (n = 14), Nocodazole (n = 13), and Nocodazole + Glyoxal (n = 13) groups of cells. Two-sided Mann– Whitney U tests (unadjusted): Untreated vs. Nocodazole, p = 7.9×10⁻⁶ (****); Nocodazole vs. Nocodazole + Glyoxal, p = 2.61×10⁻⁵ (****); Untreated vs. Glyoxal, p = 0.0275 (*). Additional contrasts (not indicated): Glyoxal vs. Nocodazole, p = 1.75×10⁻⁵; Nocodazole + Glyoxal vs. Untreated, p = 7.44×10⁻⁵; Glyoxal vs. Nocodazole + Glyoxal, p = 0.0186. Star notation: ns (p ≥ 0.05), * (p < 0.05), ** (p < 0.01), *** (p < 0.001), **** (p < 0.0001). Boxplots show the median (orange center line); the interquartile range Q1–Q3 (box); whiskers extend to the most extreme data points within 1.5× the interquartile range from the quartiles; all individual data points are overlaid. Scale bars: 10 μm.

Interestingly, even untreated cells in this dataset exhibited a detectable diffuse fraction (PolIndex = 0.65 ± 0.22), in contrast to our high-purity training datasets where filamentous integrity was near complete (PolIndex = 0.93 ± 0.04; **Supplementary** Fig. 6). This discrepancy traced back to antibody choice: an in-house conjugate with degree of labeling ∼1 yielded low background, whereas the commercial antibody used here introduced higher unspecific signal. Thus, beyond biological phenotyping, *NanTex* also functions as a quantitative quality-control tool, benchmarking labeling reagents and preparation protocols against gold-standard datasets.

Together, these data establish *NanTex* as a computational phenotyping tool that detects and quantifies structurally distinct subpopulations within a single molecular pool and reveals a glyoxal-mediated delay of nocodazole-induced depolymerization. This opens new opportunities for quantitative structural biology, from ultrastructural remodeling of the ER to mitochondrial response subtype characterization, and provides a generalizable framework for systematic benchmarking of sample preparation strategies across laboratories and users.

## Discussion

Multiplexed nanoscale imaging has long been constrained by the physics of fluorophores and detection, with spectral overlap and limited dye palettes restricting simultaneous labeling, particularly in live cells. *NanTex* reframes this problem as one of texture-based reconstruction rather than spectral separation or structural segmentation and defines a new paradigm for computational multiplexing. By exploiting nanoscale “nanotextures” as organelle fingerprints, *NanTex* achieves probabilistic demixing of single-channel images, preserving overlapping morphologies even under dense overlap conditions that segmentation-based approaches cannot resolve. Unlike recent segmentation frameworks such as Cellpose, Segment Anything, or µSAM that discretize objects, *NanTex* reconstructs overlapping organelles by leveraging nanotexture, positioning it as a complementary but fundamentally distinct class of computational multiplexing.

A central outcome of this work is the demonstration that nanotexture serves as a universal descriptor of organelle identity. Because *NanTex* requires only ∼10 representative images per organelle and modality for effective training, it can be readily adapted across systems, making it a plug-and-play solution for users who wish to extend multiplexing beyond their immediate labeling and modality constraints. Across SMLM, MINFLUX, STED, SIM, and live-cell Airyscan, *NanTex* consistently demixed identically labeled organelles ranging from cytoskeletal filaments (microtubules, actin) to vesicular systems (clathrin, endosomes, lysosomes, peroxisomes) and reticular networks (ER, mitochondria). Importantly, *NanTex* not only recovers expected structures but also uncovers hidden subdomains within single-labeled datasets, such as vesicle-like ER substructures in live cells. This ability to reveal, quantify, and track structural heterogeneity in real time makes live-cell imaging the strongest application space for *NanTex*, directly enabling computational phenotyping of organelle remodeling during cellular dynamics. Moreover, the ability to track demixed vesicular structures in ER-only datasets (**e.g. Supplementary** Fig. 4) demonstrates that *NanTex* can extend beyond visualization to provide quantitative, dynamic readouts in live-cell contexts. In nocodazole-treated HUVECs, *NanTex* separated intact microtubule filaments from diffuse depolymerized tubulin, uncovering a glyoxal-mediated phenotype not accessible with labeling-based strategies.

We note that benchmarking *NanTex* requires metrics that capture perceptual and structural fidelity. We therefore focus on SSIM and MS-SSIM, which reflect human-perceived image similarity and multi-scale preservation of structure. Segmentation-based metrics (e.g., F1/Jaccard on binarized structure masks, Manders-type assignment for channel specificity) are ill-suited, as *NanTex* does not classify discrete objects but reconstructs overlapping features. This is a critical conceptual distinction and positions *NanTex* adjacent to segmentation pipelines.

While we demonstrate the broad applicability of *NanTex*, there are caveats to be aware of (see also **Supplementary Discussion**). Out-of-focus regions and organelle heterogeneity represent two central limitations. Our blurring experiments and SMLM microtubule datasets illustrate that in areas of pronounced defocus, *NanTex* no longer assigns organelle identity, a behavior that is both expected and conceptually reassuring, as it reflects the loss of true texture information rather than spurious assignments. Similarly, organelles such as ER and mitochondria display considerable morphological diversity, ranging from fragmented to elongated networks. Importantly, this study emphasized *NanTex* to be data-efficient, with ∼10 training images per organelle, when combined with augmentation, in part to lower its adoption barrier with novice users. However, we anticipate significant performance gains with larger datasets, as well as a more robust capture of the full spectrum of organelle heterogeneity. We envision a community-wide effort to share organelle-specific SRM data that would enable training of holistic models capable of demixing ten or more organelles simultaneously.

Looking forward, *NanTex* provides a framework for significant methodological expansion. Integration with transformer-based architectures^35^ could better capture long-range structural dependencies, while adversarial (GAN)^36,46^ layers may further enhance the realism of demixed reconstructions. Beyond fluorescence SRM, the concept of texture-guided reconstruction could also extend to volumetric SRM data, correlative EM, or even (annotated) label-free modalities^45^, where structural nanotextures provide discriminative cues.

In summary, *NanTex* establishes nanotexture as a paradigm-shifting descriptor for organelle identification. By enabling label-efficient, cross-modality, and live-compatible multiplexing, it bridges computational image analysis with quantitative cell biology. Its potential for real-time phenotyping and for tracking disease-relevant remodeling positions *NanTex* as a foundation for the next generation of high-content, clinically relevant structural biology.

## Materials Availability

All data needed to evaluate the conclusions of this study are provided in the main text, supplementary information, and/or the Key Resources Table (**Supplementary Table 1**). Plasmids generated for this study have been deposited at Addgene (accession numbers listed in the Key Resources Table). No new unique reagents were generated beyond those listed.

## Data and Code Availability

All raw microscopy image files and processed source data supporting the findings of this study have been deposited at Zenodo ^51–56^ and are freely accessible. A comprehensive overview is provided in **Supplementary Table 2**. All code used for Haralick feature extraction, *NanTex* model training, inference, and evaluation (including trained network weights for all SRM modalities) is made available via **<u>GitHub</u>** and will be archived on Zenodo upon publication. Custom analysis scripts for quantitative benchmarking (SSIM, MS-SSIM, F1-score, etc.) are provided alongside training/inference pipelines in the GitHub repository. Prior to publication: *NanTex* weights are being made available to reviewers upon request. Post publication: Pretrained *NanTex* weights for SMLM, SIM, STED, and Airyscan used in this study are made available (see Zenodo/GitHub), but we emphasize that retraining may be required if imaging parameters or resolution differ substantially.

## Declaration of generative AI and AI-assisted technologies in the writing process

The authors used DeepL and ChatGPT to assist with style, grammar, and vocabulary during the preparation of this work to ensure that it is grammatically and linguistically correct. After using these tools, the authors reviewed and edited the content as needed. We, the authors, declare full responsibility for the content of the published article.

## Acknowledgements

C.F. is grateful for the support of all Franke Lab members. C.F. and C.E. especially thank Silvia Ruthardt for administrative assistance. We thank Michael Habeck and Parsa Khaleghi for fruitful discussions regarding model transferability and depolarization quantification.

C.F. is supported by the Thüringer Aufbaubank (TAB, FGR 0060 KI-supER) and the DFG via the Collaborative Research Center PolyTarget (CRC 1278, project number 316213987, project B07).

P.C. has received funding from the European Union’s Horizon 2020 research and innovation program under the Marie Skłodowska-Curie grant agreement no. 892232.

J.C.H. is supported by the Fritz-Thyssen Foundation (award number: 10.20.1.022MN), and the NCL Foundation (NCL-Stiftung).

C.K. is supported by the DFG (KA 1751/9-1).

B.T.L.V. is supported by the Thüringer Landesgraduiertenstipendien 2024 and has received funding through the Honors Programme of the Friedrich-Schiller-Universät Jena. B.T.L.V is especially grateful for all the discussions and exchange made possible by the DL@MBL 2022 initiative as well as for the funding provided therefore by the Marine Biological Laboratory and the Howard Hughes Medical Institute.

We thank the Microverse Imaging Center for providing microscope facility support. The Microverse Imaging Center is funded by the Deutsche Forschungsgemeinschaft (DFG, German Research Foundation) under Germany’s Excellence Strategy – EXC 2051 – Project-ID 39071386 and project number 316213987 (CRC 1278, project Z01). The ELYRA 7 was funded by the Free State of Thuringia with grant number 2019 FGI 0003. The LSM 980 was funded by the Free State of Thuringia with grant number 2019 FGI 0001. We further acknowledge the DFG (Instrument funding MINFLUX Jena INST 275_405_1; Instrument funding modular STED INST 1757/25-1 FUGG; GRK M-M-M: GRK 2723/1 – 2023 – ID 44711651), the State of Thuringia (TMWWDG), the Leibniz Association (Leibniz Collaborative Excellence Programme, project AMPel – project number K548/2023), and the Free State of Thuringia (TAB; AdvancedSTED / FGZ: 2018 FGI 0022; Advanced Flu-Spec / 2020 FGZ: FGI 0031) for the work on the STED and MINFLUX microscopes.

The authors also gratefully acknowledge support from the FLI Core Facilities Proteomics, and Imaging.

## Methods

### Sample Preparation for ER live-cell imaging

#### Plasmid construction

The expression construct pInd20_GFP-ATL3 was generated using the Gateway cloning system (Thermo Fisher Scientific). The entry clone pDONR221_GFP-ATL3 (kindly provided by Laura Behrendt) was recombined with the destination vector pInd20 (Addgene plasmid #44012) via an LR recombination reaction according to the manufacturer’s instructions. The resulting expression vector pInd20_GFP-ATL3 was verified by restriction digestion and sequencing.

#### Cell culture

mRFP-KDEL U2OS cells were generated as described in^21^ and were cultured in DMEM high glucose media (Sigma) supplemented with 10% heat inactivated FBS (Sigma). Cells were maintained at 37°C, 5% CO_2_ and 95% humidity in a CO_2_ incubator. Cells were seeded in 8 well chambered cover glass systems with high-precision glass coverslips (No. 1.5H, thickness 170 ± 5 µm, Thermo Fisher, cat. #155409) with 20x10^3^ cells per well. Prior to live-cell imaging of mRFP-KDEL U2OS the medium was changed to phenol-free, complete DMEM media (Thermo Fisher).

#### Generation of U2OS cell line expressing inducible GFP-ATL3

A Tet-inducible lentiviral vector (pInducer20) carrying GFP-ATL3 (pInd20_GFP-ATL3) was used to generate U2OS stable cell lines. Lentiviral particles were produced in HEK293T/LentiX cells using a three-plasmid system. Briefly, cells were plated at a density of 6.0 × 10⁶ per 10-cm dish in 10 mL growth medium (DMEM supplemented with 10% FBS and 1% Penicillin/Streptomycin). The following day, cells were co-transfected with 10 µg pInd20_GFP-ATL3, 10 µg pPAX2 (Addgene plasmid #12260), and 2.5 µg pMD2.G (Addgene plasmid #12259) using polyethyleneimine (PEI-Max, 1 mg/mL stock) in Opti-MEM (Thermo Fisher Scientific). After a 20 min complexation period, the transfection mixture was added to cells in the transfection medium (DMEM supplemented with 2% FBS, without Penicillin/Streptomycin).

Culture medium was replaced with growth medium 16–18 h post-transfection. Viral supernatants were collected at 48 h, 60 h, and 72 h post-transfection, filtered through a 0.45 µm syringe filter, aliquoted, and stored at −80 °C.

For infection, U2OS wild-type cells were plated to reach 30–40% confluency the following day and exposed to viral supernatant supplemented with 8 µg/mL protamine sulfate in a final volume of 5 mL infection medium (DMEM). In parallel, a control condition was processed identically but without viral supernatant.

Twenty-four hours post-infection, medium was replaced with complete DMEM (Thermo Fisher, cat. no. 61965-026). To select stable transductants, G418 (Geneticin) was added at a final concentration of 400 µg/mL. Medium containing G418 was renewed every 2–3 days until non-transduced control cells were fully eliminated.

#### Lysosome staining

U2OS_pInd20_GFP-ATL3-WT were plated at a density of 5.0 × 10⁴ cells per dish on 35-mm glass-bottom dishes (MatTek) in 2 mL of complete DMEM without phenol red (Thermo Fisher Scientific, cat. no. 21063-029). To induce expression of GFP-ATL3, doxycycline was added at a final concentration of 1 µg/mL. Cells were maintained under induction conditions for 48 h prior to imaging.

For lysosomal labeling, cells were incubated with LysoTracker™ Red DND-99 (Thermo Fisher Scientific) at a final concentration of 60 nM in complete DMEM. Cells were stained for 30 min at 37 °C under standard culture conditions.

### Immunofluorescence labeling of microtubules, clathrin, and ER

Cos-7 cells were grown in 8 well chambered cover glass systems with high-precision glass coverslips (No. 1.5H, thickness 170 ± 5 µm) and rinsed briefly in pre-warmed PBS (37 °C) to remove medium. Cells were permeabilized for 1–2 min using 0.3% glutaraldehyde and 0.25% Triton X-100 in cytoskeleton buffer (10 mM MES pH 6.1, 150 mM NaCl, 5 mM EGTA, 5 mM glucose, 5 mM MgCl₂), followed by fixation for 10 min in 2% glutaraldehyde in the same buffer. After two washes in PBS, residual autofluorescence was quenched with 0.1% NaBH₄ in PBS for 7 min. Cells were washed once in PBS, then blocked in 5% BSA/PBS for 30 min.

For microtubule staining, cells were incubated with rabbit anti-α-tubulin primary antibodies (2 ng/µl in blocking solution, 60 min at RT), followed by three washes (1× PBS, 2× PBS/0.1% Tween-20, 5 min each). Secondary labeling was performed with Alexa Fluor 647-conjugated goat anti-rabbit IgG (4 ng/µl, 45 min in blocking solution). Clathrin-coated pits and vesicles were stained with rabbit anti-clathrin heavy chain antibodies (2 ng/µl) and corresponding Alexa Fluor 647-conjugated goat anti-rabbit secondary antibodies (4 ng/µl). ER structures were labeled with rabbit anti-Calnexin primary antibodies in the same workflow.

After secondary labeling, cells were washed (1× PBS, 2× PBS/0.1% Tween-20) and post-fixed with 4% formaldehyde in PBS for 5 min. Final washes were performed in PBS, and samples were stored in PBS containing 0.05% sodium azide at 4 °C until imaging.

For antibody conjugation, Alexa Fluor 647-NHS was coupled to secondary goat anti-rabbit antibodies in sodium tetraborate buffer (100 mM, pH 9.5) using spin-desalting columns (40 kDa MWCO) for purification. A 2x molar dye excess was used to achieve a degree of labeling (DOL) of ∼1.1. The DOL was determined by UV–vis absorption spectroscopy.

#### HUVEC cell culture and treatment

Primary human umbilical vein endothelial cells (HUVECs) were isolated from anonymously acquired human umbilical cords according to the “Ethical principles for Medical Research Involving Human Subjects” (Declaration of Helsinki 1964) as previously described^22,43^. The protocol was approved by the Jena University Hospital Ethics Committee. The donors were informed and gave written consent. Briefly, after rinsing the cord veins with 0.9% NaCl, endothelial cells were detached with collagenase (0.01%, 3 min at 37 °C), suspended in M199/10% FCS, washed once (500 × g, 6 min) and seeded on a cell culture flask coated with 0.2% gelatin. Full growth medium (M199, 17.5% FCS, 2.5% human serum, 7.5 µg/ml ECGS, 7.5 U/ml heparin, 680 µM glutamine, 100 µM vitamin C, 100 U/ml penicillin, 100 µg/ml streptomycin) was added 24 h later. Cells were cultured until confluence and experiments usually carried out with HUVEC from the second passage.

Nocodazole-treated microtubule samples: For nocodazole experiments, HUVECs were seeded on double-coated high-precision glass coverslips (No. 1.5H, 24 mm, gelatin 1%) at a density of ∼2.2 × 10⁵ cells per coverslip in 6-well plates. Cells were cultured under standard endothelial growth conditions for 24 h before addition of glyoxal (1 mM) for 48 h.

To induce acute microtubule depolymerization, cells were treated with 10 µM nocodazole (Sigma M1404-2MG) for 5 min prior to fixation. Experimental groups included (i) untreated control cells, (ii) glyoxal-treated cells, (iii) nocodazole-treated cells, and (iv) combined GO + nocodazole treatment. Additional controls consisted of primary-only and secondary-only antibody samples.

Cells were fixed in glutaraldehyde fixation buffer for 15 min, washed with PBS, and quenched with 0.1% NaBH₄ (3 × 10 min). After permeabilization in 0.1% Triton X-100 and blocking with 3% BSA, microtubules were stained with rabbit monoclonal anti-α-tubulin antibody (ab18251, Abcam; 2 µg/ml, overnight at 4 °C) followed by Alexa Fluor 647-conjugated goat anti-rabbit secondary antibody (4 µg/ml, 1 h at room temperature). Nuclei were counterstained with Hoechst 33342 (1 µg/ml, 10 min).

### Mitochondria and peroxisomes for STED

HEK 293 (ATCC, VA, USA) and cells were maintained in a culture medium consisting of DMEM with 4500 mg glucose/L, 110 mg sodium pyruvate/L supplemented with 10% foetal calf serum, glutamine (2 mM) and penicillin–streptomycin (1%). The cells were cultured at 37 °C/8.5% CO2. Cells were grown on a #1.5 μ-Dish 35 mm. For immunolabeling, the cells were fixed with 3% formaldehyde in PBS, permeabilized for 5 min with 0,1 % Triton X-100 and blocked with 2% BSA + 5% FCS in PBS for 1 h at room temperature. Samples were incubated with primary antibodies in blocking buffer for 1 h at room temperature. To stain peroxisomes, antibodies recognizing PMP70 (Anti-PMP70 antibody (ab3421), rabbit polyclonal, Abcam, Cambridge, UK) were used. Mitochondria were labelled using an antibody against TOM20 (Tom20 Antibody (F-10): sc-17764, mouse monoclonal, Santa Cruz Biotechnology, Dallas, TX). After several washing steps, the cells were incubated for 30 min with secondary antibodies conjugated to Abberior STAR orange (rabbit) and Abberior STAR Red (mouse) (Abberior Instruments, Goettingen, Germany) diluted 1:250 in 1% BSA in PBS. After several washing steps, the slides were mounted on a drop of Mowiol (Sigma-Aldrich).

### Data Acquisition and Imaging Modalities

#### SMLM imaging

Clathrin, ER, and microtubule SMLM imaging was performed on a customized *d*STORM platform based on an inverted Olympus IX-71 microscope equipped with a 60× / 1.45 NA oil immersion objective, described in detail elsewhere^6,27^. Alexa Fluor 647 was excited with a 641 nm laser at ∼1.5-3.5 kW cm⁻² in a switching buffer consisting of 100 mM β-mercaptoethylamine (MEA), 10% (w/v) glucose, 0.5 mg ml⁻¹ glucose oxidase, and 40 µg ml⁻¹ catalase in PBS, adjusted to pH 7.4 with KOH. After a brief epi-illumination step to suppress out-of-focus fluorescence, 30,000– 50,000 frames were recorded in HiLo mode at 33 ms per frame on an Andor iXon DU-897 EMCCD camera with 128 nm optical pixel size.

Nocodazole-related datasets were acquired on a ZEISS Elyra 7 operated in *d*STORM mode, using a 63× / 1.45 NA TIRF objective and identical dye chemistry and buffer conditions as above. Frame sequences (∼20,000–50,000 frames, 25–33 ms per frame) were recorded in HiLo mode on a pco.edge sCMOS camera with 100 nm optical pixel size and processed identically in rapidSTORM^41^.

#### MINFLUX imaging

MINFLUX imaging was performed on a commercial MINFLUX system (Abberior Instruments) using identical biological samples and labeling conditions as described for SMLM, with Alexa Fluor 647 as the fluorophore. A freshly prepared imaging buffer was used containing 30 mM cysteamine (MEA), 0.5 g/L glucose oxidase (stored at –20 °C), 40 µg/mL catalase (stored at 4 °C), and 10% TRIS/Cl buffer (pH 8.0, 4 °C). For drift correction, 100 nm gold beads (fiducials) were introduced by adding 200 µL suspension into ibidi 8-well chambers for 5 min, followed by three PBS wash steps. Imaging was performed in MBM stabilization mode, optimized for long-term acquisition. Samples were pre-switched for 60–120 s at 640 nm laser wavelength and 50% laser power to reduce emitter density to the optimal range for MINFLUX localization. Subsequent imaging was carried out overnight at 10% laser power. Both 2D and 3D default imaging sequences provided by Abberior Instruments were employed without further modification.

#### SMLM Rendering and Preprocessing

Raw SMLM image stacks were processed in rapidSTORM^41^ with free FWHM fitting enabled and 500 photons lower intensity threshold. SMLM and MINFLUX localization data were rendered into grayscale 8-bit TIFF images using rapidSTORM with 10 nm/pixel resolution, intensity scaling based on photon count, and bilinear pixel-interpolation. Localization files were converted from proprietary formats (.smlm, .msr, etc.) into plain-text files readable by rapidSTORM using custom scripts if necessary.

Rendered images were cropped and converted to uniform resolution and scale. For MINFLUX data, additional histogram equalization and morphological opening (2-pixel kernel) were applied to suppress depth-projected artifacts.

#### SIM imaging

SIM imaging was performed on a ZEISS Elyra 7 system (laser module: 000000-0239-500) equipped with four laser lines (405, 488, 561, 641 nm) and a mercury vapor lamp. A Plan-Apochromat 63×/1.4 oil immersion objective was used in combination with the built-in Optovar (1.6×), yielding an effective pixel size of 62 nm at the detector. For comparative *d*STORM measurements, a 63×/1.45 NA oil immersion TIRF objective was employed. A quad-band dichroic beamsplitter and emission filter (LBF 405/488/561/642) was used for all acquisitions.

Each SIM dataset was acquired in lattice SIM mode as a z-stack with a physical step size of 110 nm, where each plane consisted of 13 phase-shifted raw images. 3D datasets were recorded to optimize optical sectioning and suppress out-of-focus contributions, but only 2D SIM reconstructions were used for downstream analysis. Exposure time per phase was 250 ms, and optical gratings were selected according to the system’s default settings. SIM reconstruction was carried out in ZEN Black (Zeiss) with the internal reconstruction tool using the "precise" quality setting, no baseline cut, and default Wiener filtering.

#### STED imaging

STED imaging was performed on an Abberior STEDYCON system (Abberior Instruments GmbH) mounted on an Olympus IX83 inverted microscope equipped with a UPlanXApo 100×/1.45 NA oil immersion objective. Images were acquired with an effective pixel size of 20 nm, a pixel dwell time of 10 µs, and 3–10 line accumulations depending on the size of the region of interest. Mitochondria were labeled with Abberior STAR ORANGE and excited with a pulsed 561 nm laser (0.5–3.2 µW, adjusted to labeling density), while peroxisomes were labeled with Abberior STAR RED and excited with a pulsed 640 nm laser (16–33 µW). In both cases, fluorescence depletion was achieved with a pulsed 775 nm fiber laser operated at ∼ 225 mW in the sample plane. Reported laser powers correspond to mean values at the specimen plane; values were approximated based on calibrated power measurements taken on different experimental days and adjusted with internal correction factors for optical losses. Emission signals were split between two avalanche photodiode detectors (SPCM-AQRH, Excelitas Technologies): the STAR ORANGE channel was collected between 580–630 nm, and the STAR RED channel between 650–700 nm. Time-gated detection was applied between 1–7 ns after excitation. All acquisitions were controlled using STEDYCON Smart Control software (version 7.1.53, Abberior Instruments).

#### Airyscan live-cell imaging

Airyscan live-cell imaging was performed on a Zeiss LSM 980 confocal microscope equipped with an Airyscan2 detector in Fast 2D mode. Cells were maintained at 37 °C and 5% CO₂ in a stage-top incubation chamber (PeCon). Images were acquired using a Plan-Apochromat 63×/1.4 oil immersion objective. GFP-ATL3 was excited at 488 nm and detected at 523 nm, while LysoTracker Red was excited at 561 nm and detected at 579 nm; mRFP was excited at 590 nm the emission was captured from 612 nm. Sequential image series were collected with a pixel size of 43 nm selecting a region of interest of 26.48 × 26.48 µm, resulting in a frame time interval of ∼ 260 ms. Images were optionally deconvolved with the Airyscan processing tool of the ZEN Blue software (v3.10).

### Haralick Feature Extraction and Classical Clustering

#### Feature computation

Classical textural descriptors were generated using the *Mahotas* Python library (v1.4.13)^38^. For each pixel, a set of 13 Haralick features were extracted from gray-level co-occurrence matrices (GLCMs), including angular second moment, contrast, correlation, variance, inverse difference moment, sum average, sum variance, sum entropy, entropy, difference variance, difference entropy, information measures of correlation, and maximal correlation coefficient. To capture local heterogeneity, both the mean and peak-to-peak range of each Haralick feature were calculated within a 7×7 sliding window. This resulted in a 26-dimensional feature vector per pixel.

#### Normalization

All feature values were scaled to the range [0,1] by min–max normalization, applied independently per feature. Scaling ensured comparability across different samples and avoided dominance of high-variance features. Feature maps were border-trimmed during sliding-window calculation to prevent artifacts at image edges.

#### F1-Score

Haralick feature maps were generated from computational superimpositions of two mono-structural images. These input images were then masked to isolate the regions of interest. To evaluate segmentation performance, we compared the nonzero pixels in the overlay of the original images with the nonzero pixels in the thresholded feature maps. This comparison was performed across multiple threshold levels. The performance of each threshold was quantified using the F1-score as implemented in *<u>scikit-learn</u>* (v1.2.1), with the highest score being returned for a comparison. As semantic segmentation is a binary decision on the pixel level (either Microtubules or LNP), regions of overlap were considered to be belonging to the more present structure, in this case Microtubules.

#### Clustering

Unsupervised clustering was performed on the 26-dimensional feature stack using k-means (k = 5) as implemented in *<u>scikit-learn</u>* (v1.1.3). Each pixel was assigned to one of five clusters by minimizing squared Euclidean distance in feature space. This procedure yielded cluster maps representing regions of locally similar nanotexture.

#### Evaluation

While Haralick clustering proved useful for feature exploration and highlighted major differences in nanotexture between organelles, it was insufficient for precise multiplexing in regions of structural overlap. Specifically, cluster boundaries frequently fragmented continuous structures (e.g., filamentous microtubules) and failed to preserve co-localized signals. These observations motivated the development of *NanTex*, which extends beyond classical feature-driven segmentation to probabilistic demixing of overlapping structures.

### NanTex

#### Computational superimposition and first level data augmentation

To generate train, test and validation data, mono-structural images were rescaled [0, 255], resized (padded or cut), rotated against each other (0°, 90°, 180°, 270°), and overlaid summatively. Permuting over all combinations of samples per structure and rotations, the amount of available data had been increased substantially while creating different structural aggregates.

#### Data preprocessing and second level augmentation

All input fluorescence images were preprocessed by standardizing each patch to zero mean and unit variance. Ground-truth channels corresponding to individual organelles were min–max normalized to the range [0,1]. Basic augmentation was applied using the *<u>Albumentations</u>* library^37,38^ to increase model robustness to photophysical variability and imaging artifacts. Augmentations included: randomized patch cutouts (256x256), randomly applied horizontal and vertical flips, and Median blur (kernel size in [3,5]. All data processing involved has been performed on an *AMD Ryzen Threadripper 2920X 12-Core CPU* leveraging a custom implementation of the *<u>ray</u>* library to realize multiprocessing.

#### Network architecture

*NanTex* employs a UNet–style convolutional encoder–decoder with four resolution levels. Each level consists of two 3×3 convolutions followed by batch normalization and ReLU activation. Downsampling is implemented via 2×2 max pooling, while upsampling uses transposed convolutions. Skip connections ensure retention of fine spatial information. The network processes 32 feature maps per patch. The demixing head is a 1×1 convolutional layer with linear activation, producing a grayscale output for each channel (256×256 each), corresponding to the predicted organelle structures.

#### Training protocol

Training was performed in *<u>PyTorch</u>* (v1.12)^39^ on an NVIDIA RTX A5000 GPU. The Adam optimizer was used with learning rate 1×10−5, β1=0.9, β2=0.99, and ε=1×10−8. The loss function was defined as mean squared error (MSE) between predicted and ground-truth images. Variable batch sizes (e.g. 16, 32 patches) were used. Training typically converged after ∼80–120 epochs of varying sizes (e.g. 16, 32, 64 cycles), monitored by validation MSE and SSIM.

#### Prediction and post-processing

Multi-structural images were subsequently loaded, resized (to prevent shape mismatches), standardized to zero-mean-unit-variance, and divided into patches (usually 256x256 px), before being passed through the model sequentially, skipping empty patches to increase execution speed and prevent hallucinations. Predicted patches were reassembled into full-size images using a sliding window with stride = 256 px (no overlap). Background correction was applied by histogram-based binning, where the five most frequent intensity values per channel were zeroed to suppress background contributions.

#### Quantitative evaluation with SSIM and MS-SSIM

To benchmark *NanTex* reconstructions against ground-truth (GT) channels, we used the *<u>pytorch-</u> <u>msssim</u>*implementation of the Structural Similarity Index (*SSIM*) and its multi-scale extension (*MS-SSIM*), two perception-aligned image quality metrics that compare local luminance, contrast, and structure rather than raw pixel errors. SSIM is computed on small windows and combines three terms, luminance, contrast, and structure, into a single score in [−1, 1] (practically [0, 1] after intensity normalization). MS-SSIM extends SSIM across image pyramids, weighting coarse and fine scales to better reflect human/perceptual judgment and robustness to small misregistrations^29–31^. We report both because SSIM is highly sensitive to high-frequency losses (e.g., blur), while MS-SSIM more faithfully summarizes fidelity across spatial scales (useful when nanotexture is preserved but slightly smoothed). For completeness, PSNR was also computed as an internal quality control metric but not used for decision-making, as it correlates weakly with perceived quality compared to SSIM-type metrics.

#### Computation

All images were 8-bit or normalized to [0, 255]. SSIM used a Gaussian window (11×11, σ = 1.5), constants C1 = (0.01L)², C2 = (0.03L)² with dynamic range L = 1, and “reflect” boundary handling (scikit-image, v0.22; parameters per default)^40^. MS-SSIM used 5 scales with down-sampling by 2 and the original weighting vector [0.0448, 0.2856, 0.3001, 0.2363, 0.1333]^30^. Metrics were computed (i) globally per full field of view and (ii) locally on 256×256 patches/ROIs (reported as mean ± s.d. across patches). All scores are reported per GT/predicted organelle channel pair.

#### Controls and caveats

i. SSIM can be deteriorated by large uniform backgrounds.
ii. Small lateral misregistrations depress SSIM; MS-SSIM mitigates this but does not replace proper alignment.
iii. Two unrelated images with similar low variance can yield mediocre SSIM; our negative controls (unrelated organelles, natural images, voids) produced low SSIM/MS-SSIM, validating specificity (Supplementary Fig. 2).
iv. Because SSIM-type metrics remain intensity-aware, we also report intensity-agnostic metrics elsewhere when the biological question concerns presence/absence rather than brightness (e.g., computational phenotyping).

#### Modality-specific considerations

Notably, in STED and Airyscan datasets, single-scale SSIM values were reduced due to the inherently lower SNR and contrast of these modalities, yet MS-SSIM remained high (0.86–0.90 for STED; > 0.74 for Airyscan), accurately reflecting preservation of large-scale organelle morphology. This highlights that multi-scale metrics are more appropriate for benchmarking *NanTex* in lower-SNR imaging regimes where local contrast is degraded but global structures are faithfully demixed.

#### Reporting

For each modality and organelle type we report global and patch-wise SSIM/MS-SSIM (main text and figure panels), and we visualize per-pixel SSIM heatmaps to localize discrepancies where meaningful. When summarizing across many fields, we present distributions (box plots) and the median with interquartile range.

#### Interpretation (rule-of-thumb for *NanTex* SRM demixing)

Absolute thresholds depend on modality/SNR. In our experience, MS-SSIM is typically slightly higher than SSIM when large-scale morphology is correct but fine detail is modestly smoothed; the converse (MS-SSIM ≤ SSIM) flags scale-dependent artifacts (e.g., high-frequency hallucinations). We therefore use both metrics jointly: high MS-SSIM with high SSIM indicates faithful demixing; divergence between the two prompts visual inspection or complementary, intensity-agnostic measures if applicable. **Table 1** provides an overview consistent with qualitative meaning in our datasets and controls (also see **Supplementary** Fig. 2)

### Single Particle Tracking of Demixed Vesicles

Demixed time-lapse image stacks were analyzed using TrackMate (v7.14.0)^49^. Vesicles were detected with the LoG (Laplacian-of-Gaussian, 15 pixel object diameter, quality threshold = 0.5) detector with sub-pixel localization enabled and the median filter option active to suppress noise. Tracks were built with the advanced Kalmann Tracker (search radius 20 px, max frame gap 2 frames). Mean-squared displacement (MSD) curves were computed per track from the exported coordinates and processed with custom Python scripts as described in detail in ^50^.

### Computational Phenotyping of Microtubule Depolymerization

Computational phenotyping of microtubule morphology subtypes was performed on *d*STORM images of untreated, glyoxal-treated, nocodazole-treated, and combined glyoxal/nocodazole-treated HUVEC cells. The *NanTex* model trained on untreated microtubule SMLM, clathrin and ER data was applied to demix each image into two channels: (i) intact microtubules (microtubule channel) and (ii) diffuse tubulin (combined other channels). For each cell, the number of positive pixels per channel was determined after thresholding at the 95th percentile of background intensity distribution to exclude noise. The *polymerization index* was then defined straight forwardly as

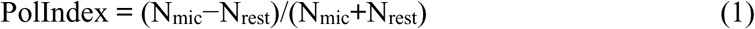

where N_mic_ denotes the number of pixels assigned to the intact microtubule channel and N_rest_ the number of pixels assigned to the diffuse tubulin channel. This metric provides a normalized, cell-wise readout ranging from –1 (entirely depolymerized) to +1 (entirely filamentous). Values closer to zero correspond to mixed phenotypes or unspecific backgrounds.

Single-cell *polymerization index* values were compiled across conditions to assess treatment-dependent phenotypes. Glyoxal treatment alone yielded intermediate values, whereas nocodazole exposure produced a strong shift toward negative indices, consistent with microtubule depolymerization. The combined treatment condition enabled assessment of potential additive or synergistic effects.

Statistical analysis was performed on the per-cell polymerization index distributions across conditions. Group comparisons were evaluated using Welch’s *t*-tests alongside non-parametric Mann–Whitney *U* tests to account for small sample sizes and potential variance heterogeneity. Effect sizes were additionally reported (Cohen’s *d*, rank-biserial correlation) to capture the magnitude of differences independent of sample size. P-values < 0.05 were considered significant. Given the limited sample sizes for these proof-of-concept experiments (n = 3–5 cells per condition), statistical interpretation relied not only on *p*-values but also on effect size measures, which provide a more robust estimate of biological relevance under small-*n* conditions. Welch’s *t*-tests were used for group comparisons allowing unequal variances, and Mann–Whitney *U* tests served as a non-parametric validation. Effect sizes were reported as Cohen’s *d* (parametric) and rank-biserial correlation (non-parametric). This combined approach ensures that both statistical significance and the magnitude of observed effects are transparently communicated, with effect sizes serving as the primary criterion where statistical power was limited.

## Supplementary Figures

**Supplementary** Figure 1. Illustration of SSIM and MS-SSIM specificity as controls for *NanTex* evaluation.

**Supplementary** Figure 2. Resolution-dependence of NanTex performance.

**Supplementary** Figure 3. Suboptimal *NanTex* performance with AF647 STED data.

**Supplementary** Figure 4. *NanTex* demixing and single particle tracking of live-cell Airyscan datasets.

**Supplementary** Figure 5. Validation of *NanTex* recognition specificity using reduced localization density.

**Supplementary** Figure 6. High-purity microtubule *d*STORM datasets used as reference for quality control.

## Supplementary Videos

**Supplementary Video 1.** Live-cell Airyscan: ER–lysosome dynamics demixed by *NanTex*.

**Supplementary Video 2.** *NanTex* demixing of ER-only super-resolution Airyscan live-cell data.

