## Supplementary Information for "NanTex enables computational multiplexing and phenotyping of organelles across super-resolution modalities"

### Supplementary Discussion: Current Limitations and Outlook

While *NanTex* provides a powerful framework for computational multiplexing across diverse SRM modalities, several limitations must be considered when interpreting and extending our findings.

1. **Dependence on training data quality**

   *NanTex* performance is strongly influenced by the resolution and labeling quality of the training datasets. Networks trained on clean SMLM data generalized successfully to MINFLUX but deteriorated on resolution-degraded inputs (Supplementary Fig. X). Thus, *NanTex* is not resolution-invariant, and high-quality, representative datasets remain essential for robust performance. This is not surprising, as suboptimal labelling will always lessen the information content and validity of imaging data, thus *NanTex* follows the common rule: “Garbage in, garbage out!”

2. **Contrast sensitivity across modalities**

   The method relies on sufficient structural contrast to extract nanotextural features. This was evident in STED experiments with AF647, a dye known to perform suboptimally under depletion conditions. In contrast, optimized dye combinations (AbStar Red/Orange) produced reliable multiplexing results. Careful fluorophore and protocol selection are therefore critical for successful application in new imaging systems. Referring to the point above, *NanTex* trained on an organelle with a particular dye might not perform as expected when tasked with data from the same organelle, but another dye - this will be especially true for SRM modalities that heavily rely on photophysical properties, i.e. SMLM and STED.

3. **Influence of rendering parameters on nanotexture extraction**

   A further methodological consideration lies in the rendering of SMLM localizations into images. In the early development of SMLM, it was recognized that rendering strategies (e.g., Gaussian vs. histogram rendering, kernel widths, pixel sizes, intensity normalization etc.) can substantially influence the apparent structural detail of reconstructed images. Since *NanTex* relies on nanotextural cues, these rendering choices directly affect the fidelity of extracted features. For example, overly smooth Gaussian kernels may obscure fine structural texture, whereas undersampling may fragment continuous filaments or vesicles. While our study employed standardized rendering parameters to ensure reproducibility, it is conceivable that adapted or data-specific rendering approaches could further enhance nanotexture preservation and thereby improve *NanTex* performance. Future pipelines may even bypass classical rendering entirely, operating directly on localization coordinates to avoid representation bias. Rendering considerations are also valid for other SRM modalities and will be studied in more detail in the future.

4. **Interpretative boundaries of demixing**

*NanTex* separates structures based on textural features rather than molecular identity. While this enables discovery of previously unresolvable subpopulations (e.g., ER-mediated vesicles), it can also generate biologically ambiguous channels if structural differences are subtle or non-specific (e.g. SMLM data of ER and clathrin both exhibited indistinguishable diffuse fractions). While this effect is negligible in high quality, high contrast data, it becomes relevant when working with data of lesser quality. In these cases, independent validation, for example with orthogonal molecular markers or correlative EM, remains essential for biological interpretation. These could also be integrated into the training of the network as ground truth references.

5. **Breadth versus throughput**

Our validation encompassed a representative set of organelles (microtubules, actin, clathrin, ER, endosomes, lysosomes, mitochondria, peroxisomes). While *NanTex* conceptually generalizes, each novel organelle or structure requires sufficient high-quality training data. Rare or transient morphologies may remain underrepresented until captured in larger datasets.

6. **Data efficiency versus model quality**

In our presented work we deliberately chose to showcase *NanTex* data efficiency by relying on heavy data augmentation, so that ~ 10 input images per organelle and imaging mode were sufficient to train and test the models. This was to enhance the applicability of our approach and lower the threshold for non-experts that might not have the infrastructure of acquiring large numbers of data. However, it seems reasonable that increasing the amount of high-quality training data will enhance *NanTex* multiplexing and phenotyping performance. We suggest a SRM community effort to collect larger high quality data sets for training and validation - see below.

7. **Computational requirements**

Although inference is rapid, training *NanTex* models requires substantial GPU resources, particularly when retraining for new modalities or organelle classes. Broader adoption may thus benefit from pre-trained community models and shared benchmarking datasets.

8. **Quantitative benchmarking**

Our validation employed SSIM and MS-SSIM as image-based similarity metrics, which primarily capture structural fidelity but not biological correctness. Future benchmarking against orthogonal ground truths (e.g., correlative EM, functional readouts) will be essential to firmly establish *NanTex* as a quantitative standard for structural phenotyping.

#### Outlook

Despite these limitations, *NanTex* addresses a long-standing bottleneck in SRM: spectral and temporal constraints in multiplexed imaging. By leveraging structural texture rather than spectral diversity, *NanTex* opens new avenues for studying highly overlapping organelles, unlabeled structural subtypes, and dynamic processes in live cells. Integration with spectroscopy-based multiplexing, correlative EM, and standardized benchmarking pipelines may further expand its utility. Collectively, *NanTex* offers a first step toward computationally unlocking multiplexed nanoscale biology at scale, complementing both existing labeling strategies and future advances in probe chemistry. *NanTex* was intentionally demonstrated with minimal training data and heavy augmentation to facilitate accessibility for new users. Larger, more diverse datasets are expected to markedly improve fidelity.

**We therefore advocate for a community-driven repository of organelle-specific SRM datasets to train universal *NanTex* models capable of multiplexing many organelles simultaneously.**

Second, while our focus here was fluorescence-based SRM, the same approach could be extended to correlative EM or label-free microscopy if sufficiently annotated training data were available. Finally, future iterations of *NanTex* could incorporate transformer backbones to model global context and GAN-based modules for realistic reconstruction, offering a path toward even more robust and generalizable multiplexing. Looking ahead, integration with community-driven SRM datasets, akin to an ImageNet-style resource for nanoscale imaging, could accelerate the development of universal *NanTex* models capable of demixing a broad spectrum of organelles simultaneously.

**Supplementary Table 1 | Key Resources Table.** Reagents, cell lines, plasmids, chemicals, software tools, and microscopy systems used in this study. Plasmids generated here have been deposited to Addgene (see identifiers). All microscopy raw data and *NanTex* model weights are deposited to Zenodo. Custom code for feature extraction and model training is available on GitHub.

| REAGENT or RESOURCE | SOURCE | IDENTIFIER |
|---|---|---|
| CELL LINES | | |
| HEK293T | ATCC | CRL-3216 |
| Cos-7 | Cell Lines Service GmbH | #605470 |
| U2OS_pInd20_GFP-ATL3-WT | This study / Gift from collaborator | Available upon request |
| HUVEC (primary) | Anonymously acquired umbilical cords | |
| mRFP-KDEL U2OS | This study / Gift from collaborator | Available upon request |
| ANTIBODIES | | |
| Mouse anti-α-tubulin | Sigma-Aldrich | T5168 |
| Rabbit anti-α-tubulin | Abcam | ab18251 |
| Rabbit anti-clathrin heavy chain | Abcam | ab21679 |
| Rabbit anti-Calnexin | Abcam | ab22595 |
| Goat anti-mouse Alexa Fluor 647 | Invitrogen | A-21236 |

| | | |
|---|---|---|
| Goat anti-rabbit Alexa Fluor 647 | Invitrogen | A-21245 |
| Goat anti-rabbit IgG | Invitrogen | #31212 |
| Goat anti-rabbit Abberior STAR Red | Abberior | STRED-1002 |
| Goat anti-rabbit Abberior STAR Orange | Abberior | STORANGE-1001 |
| CHEMICALS AND REAGENTS | | |
| Alexa Fluor 647 NHS-Ester | Life Technologies | A-20106 |
| Nocodazole | Sigma-Aldrich | M1404-2MG |
| Glyoxal | Sigma-Aldrich | 50649 |
| β-mercaptoethylamine (MEA) | Sigma-Aldrich | M6500 |
| Glucose oxidase | Sigma-Aldrich | G2133 |
| Catalase | Sigma-Aldrich | C9322 |
| LysoTracker™ Red DND-99 | Thermo Fisher Scientific | L7528 |
| Hoechst 33342 | Thermo Fisher Scientific | H3570 |
| sodium tetraborate | Fulka | 71999 |

| | | |
|---|---|---|
| DMEM medium | Sigma | D6429 |
| DMEM complete medium | Thermo Fisher Scientific | 11054020 |
| Fetal bovine serum (FBS, heat inactivated) | Sigma | F7524 |
| PLASMIDS | | |
| pInd20_GFP-ATL3-WT | This study, as in[21] | Addgene #44012 |
| MICROSCOPY HARDWARE | | |
| Zeiss Elyra 7 lattice SIM | Carl Zeiss Microscopy | System ID available upon request |
| Zeiss LSM 980 Airyscan | Carl Zeiss Microscopy | System ID available upon request |
| Leica STED Infinity Line | Leica Microsystems | System ID available upon request |
| Abberior STEDYcon | Abberior Instruments | System ID available upon request |
| Custom-built *d*STORM (based on Olympus IX-71) | This study / as in [6,27] | — |
| Abberior MINFLUX (commercial) | Abberior Instruments | System ID available upon request |
| SOFTWARE AND ALGORITHMS | | |
| Zen Black / Zen Blue | Carl Zeiss Microscopy | https://www.zeiss.com/microscopy |
| rapidSTORM | Wolter et al.[41] | https://github.com/super-resolution/rapidSTORM |
| scikit-image (v0.22) | van der Walt et al.[40] | https://scikit-image.org |

| | | |
|---|---|---|
| Albumentations (v1.3) | Buslaev et al.[37] | https://albumentations.ai |
| PyTorch (v1.12) | Paszke et al.[39] | https://pytorch.org |
| *NanTex* (this study) | This study, GitHub | https://github.com/Zatyrus/NanTex |
| Haralick feature extraction scripts | This study, GitHub | https://github.com/Zatyrus/NanTex |
| DATA REPOSITORIES | | |
| Microscopy raw data & training datasets | This study, Zenodo | [51-56] |
| *NanTex* model weights (per modality) | This study, Zenodo | **Prior publication:** available for reviewers upon request **Post publication: open source** available via Zenodo |

**Supplementary Table 2 | *NanTex* data sets. |** Overview and metadata of the computationally superimposed image stacks used in training, validation and testing of *NanTex* per SRM modality. $N_x$ corresponds to the number of mono-structural samples used per organelle and modality.

| Modality | Structures | $N_{train}$ | $N_{val}$ | $N_{test}$ | Source / Microscope |
|---|---|---|---|---|---|
| SMLM I | Microtubules | 13 | 2 | 4 | Shareloc |
| | LNPs | 4 | 1 | 2 | Shareloc |
| | Actin | 3 | 1 | 1 | Shareloc |
| SMLM II | Microtubules | 8 | 1 | 5 | Olympus IX-71 |
| | ER | 4 | 1 | 3 | |
| | Clathrin | 6 | 1 | 2 | |
| SIM | Microtubules | 15 | 5 | 5 | ZEISS Elyra 7 |
| | Clathrin | 15 | 5 | 5 | |
| STED I | Microtubules | 10 | 2 | 2 | Leica Infinity Line |
| | Clathrin | 10 | 2 | 2 | |
| STED II | Peroxisomes | 10 | 4 | 2 | Abberior STEDYCON |
| | Mitochondria | 10 | 4 | 2 | |
| Airyscan | Lysosomes | 20 | 5 | 20 | Zeiss LSM 880 Airyscan |
| | ER | 20 | 5 | 20 | |

### Supplementary Figures

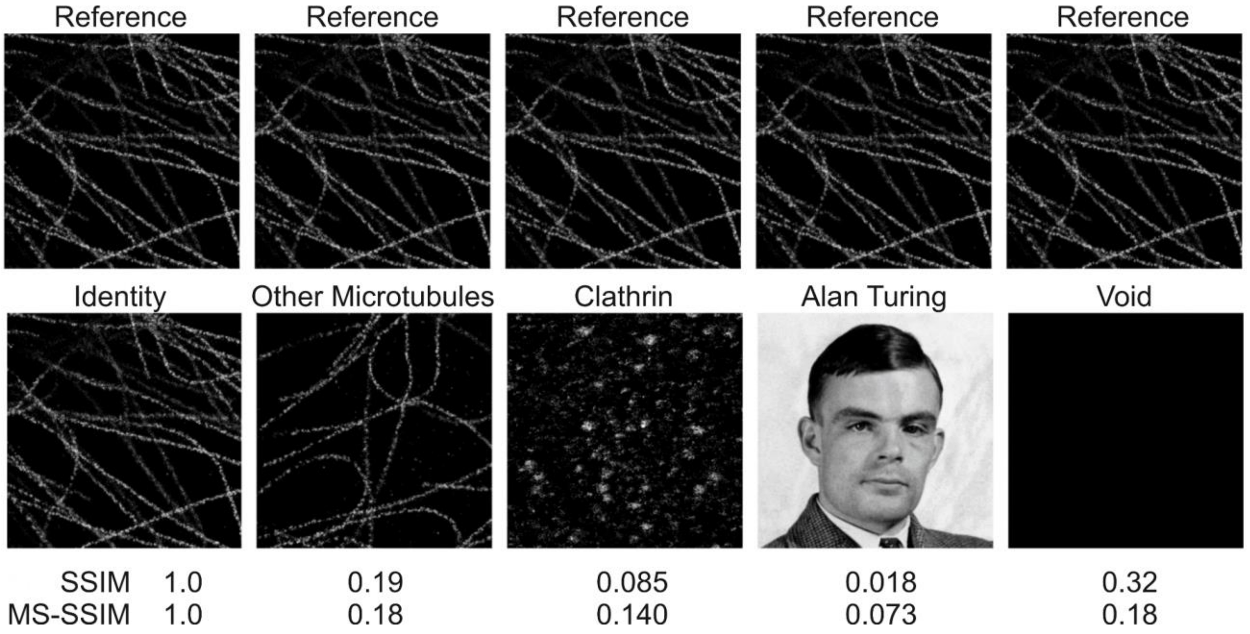


**Supplementary Figure 1 | Illustration of SSIM and MS-SSIM specificity as controls for *NanTex* evaluation.** Representative *d*STORM microtubule reference image (top row) was compared against various test images (bottom row), including the identical microtubule image (Identity), an unrelated microtubule image (Other Microtubules), clathrin, a non-microscopy control (portrait of Alan Turing), and an empty image (Void). Structural similarity (SSIM) and multi-scale SSIM (MS-SSIM) were computed for each comparison and are shown below. Only the identical image yields perfect similarity (SSIM = 1.0, MS-SSIM = 1.0), whereas all unrelated images result in low values. This demonstrates that SSIM-based metrics are highly specific for structural agreement and do not falsely recognize unrelated content, thereby validating their use for quantitative benchmarking of *NanTex* reconstructions.

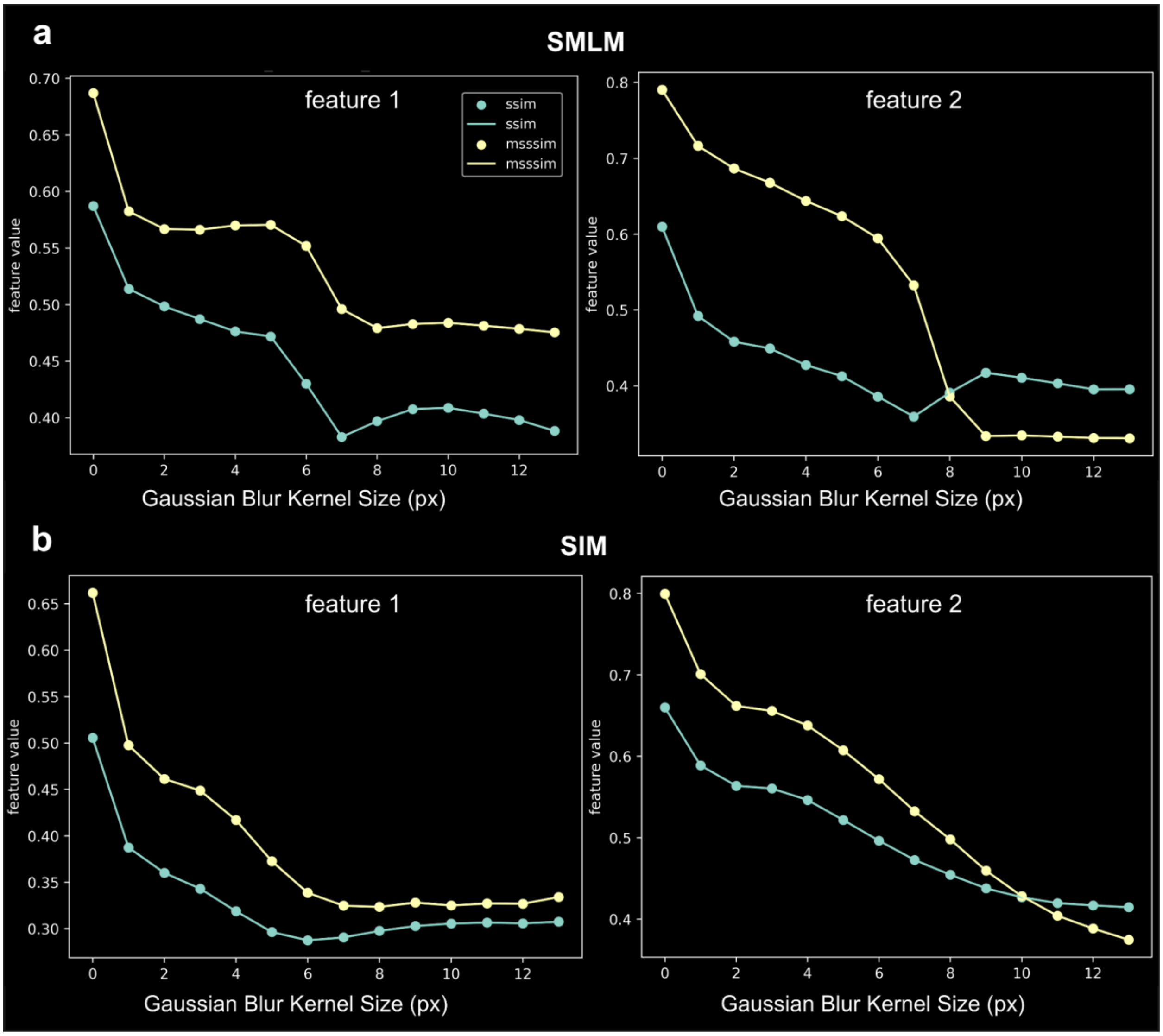


**Supplementary Figure 2 | Resolution-dependence of *NanTex* performance.** To systematically assess how resolution impacts *NanTex* fidelity, high-quality ***d***STORM (**a**) and SIM (**b**) microtubule datasets were progressively degraded by Gaussian blurring of increasing kernel size (x-axis). Demixed channels were compared against the corresponding ground truth using SSIM (cyan) and MS-SSIM (yellow). Both modalities showed a monotonic decline in fidelity with increasing blur, with MS-SSIM generally more robust than SSIM to small-scale degradation. Notably, SSIM/MS-SSIM values dropped sharply instantly, indicating a resolution threshold below which nanotexture features are no longer reliably preserved compared to training data. These results validate that *NanTex* recognition depends on genuine structural detail rather than global intensity cues and highlight the necessity of sufficient resolution for accurate computational multiplexing.

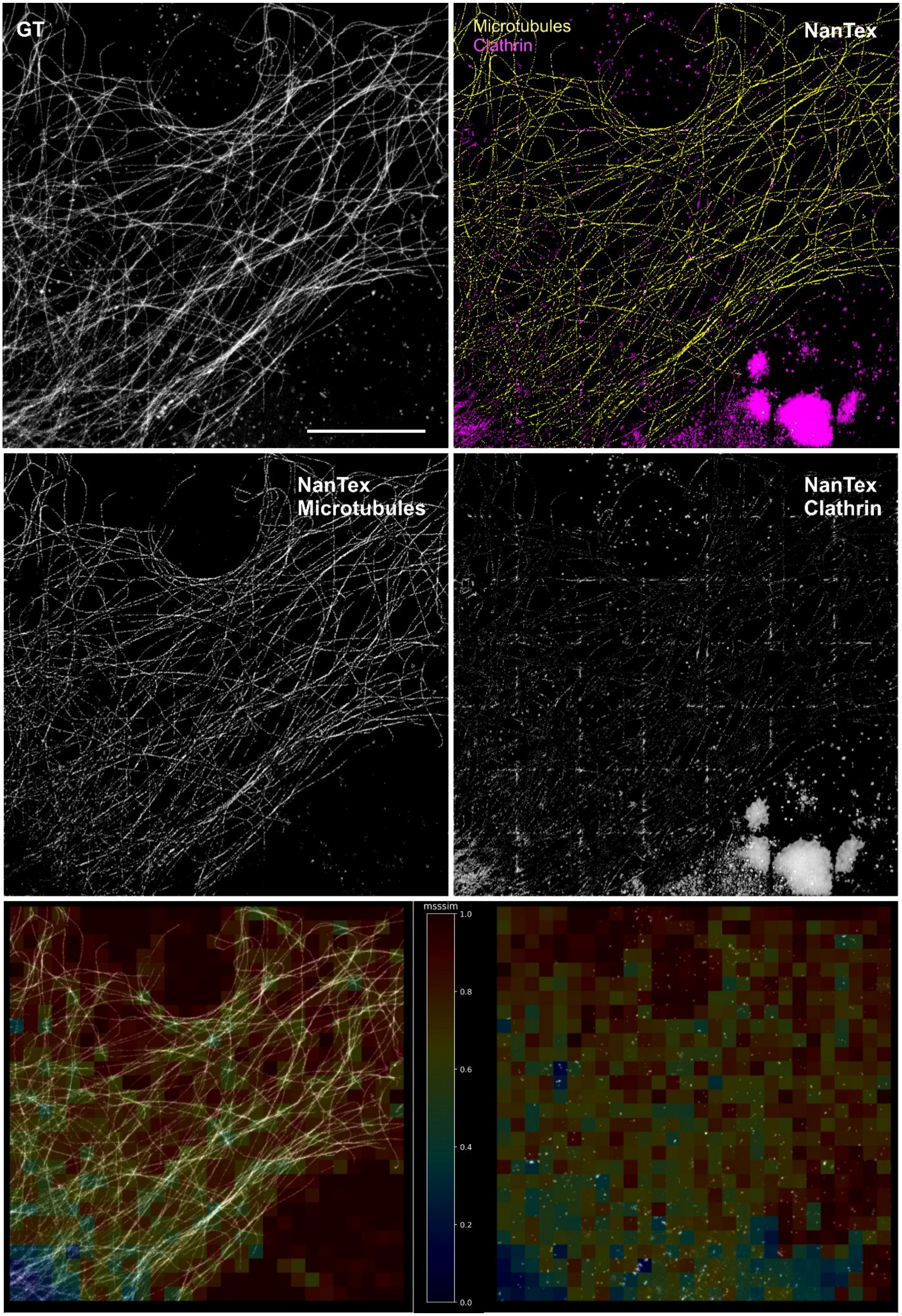
GT
Microtubules
Clathrin
NanTex
NanTex
Microtubules
NanTex
Clathrin
msssim
1.0
0.8
0.6
0.4
0.2
0.0

**Supplementary Figure 3 | Suboptimal *NanTex* performance with AF647 STED data.** *Upper left:* Representative ground truth STED image of AF647-labeled clathrin and microtubules, acquired under depletion conditions suboptimal for this fluorophore. *Upper right: NanTex* multiplexing results, shown in false colors (yellow: microtubules, magenta: clathrin). *Middle: NanTex* results and corresponding MS-SSIM heatmaps (*bottom*). While microtubules are recognized with reasonable fidelity, the clathrin channel shows several artefacts, including false positive large ghost structures and an overlayed checkered pattern, stemming from insufficient contrast in the source images. Scale bar: 10 µm.

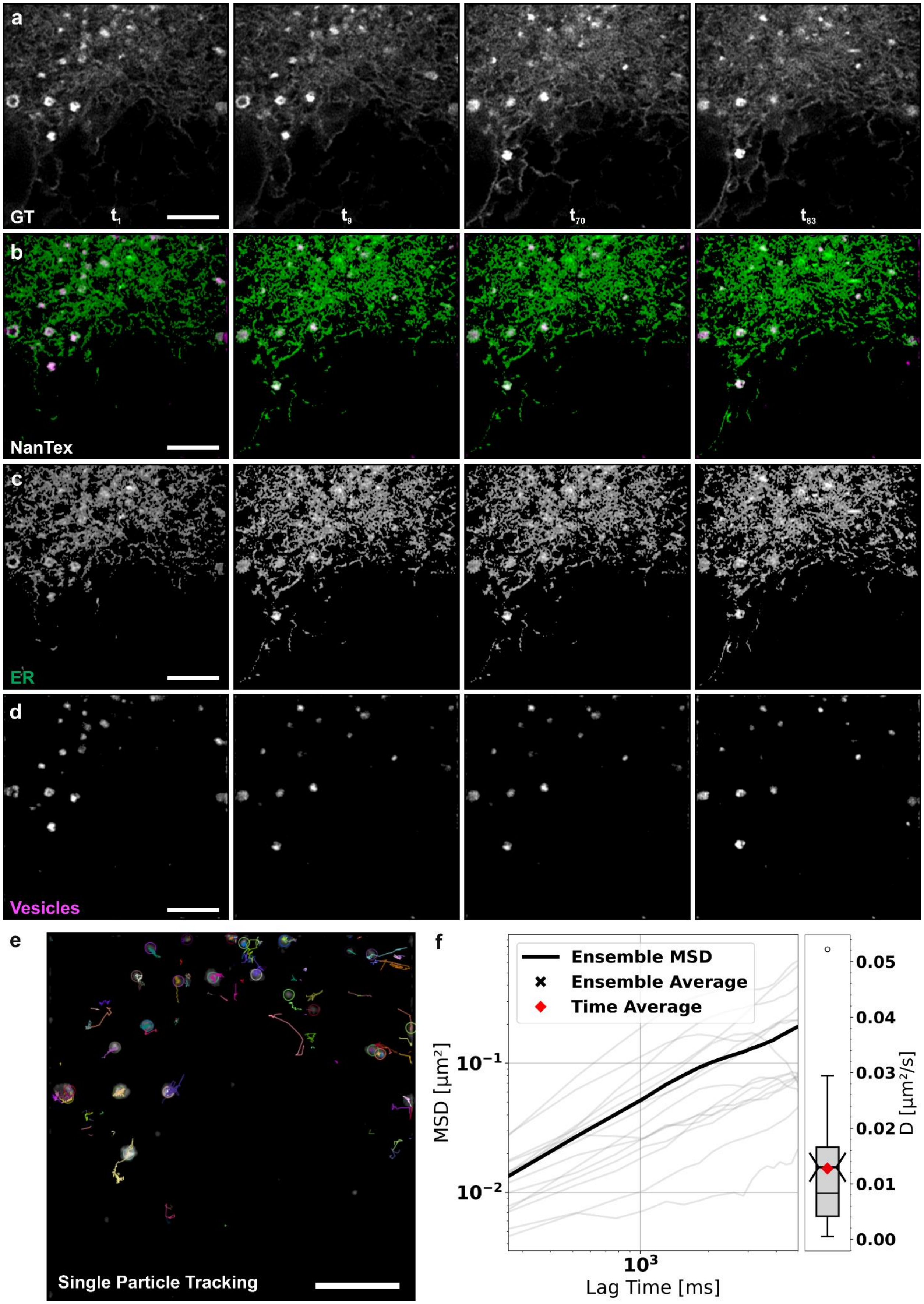
a
GT
$t_1$
$t_9$
$t_{70}$
$t_{83}$
b
NanTex
c
ER
d
Vesicles
e
Single Particle Tracking
f
Ensemble MSD
Ensemble Average
Time Average
MSD [μm²]
Lag Time [ms]
D [μm²/s]

138

**Supplementary Figure 4 | *NanTex* demixing and single particle tracking of live-cell Airyscan datasets**, where only the ER is labeled (mRFP-KDEL U2OS) with representative timepoints (entire time course shown in **Supplementary video 2**). (**a**): ground truth grey-scale data, contrast / dynamic range enhanced for visibility. (**b**): *NanTex* results in false colors showing ER in green and vesicle-like ER substructures in magenta. (**c,d**) Panels show single channel *NanTex* results in greyscale. (**e**) *NanTex*-enabled single-particle tracking of demixed vesicle-like structures using TrackMate[49]. (**f**) Mean-squared displacement (MSD) analysis[50] of individual vesicle tracks with more than 29 frames (grey) and the ensemble average (black) confirm dynamic behavior of ER-associated vesicles. We display the distribution of diffusion coefficients (time-average: 0.013±0.07 $\mu m^2/s$ (red diamond) and ensemble-average: 0.013±0.01 $\mu m^2/s$ (black cross)) extracted from the MSD as a boxplot (center line, median; red diamond, mean; box limits, first to third quartile; whiskers, 1.5x interquartile range; points, fliers) to the right. This highlights how *NanTex* makes quantitative vesicle tracking feasible from single-channel ER datasets. Scale bars 10 µm.

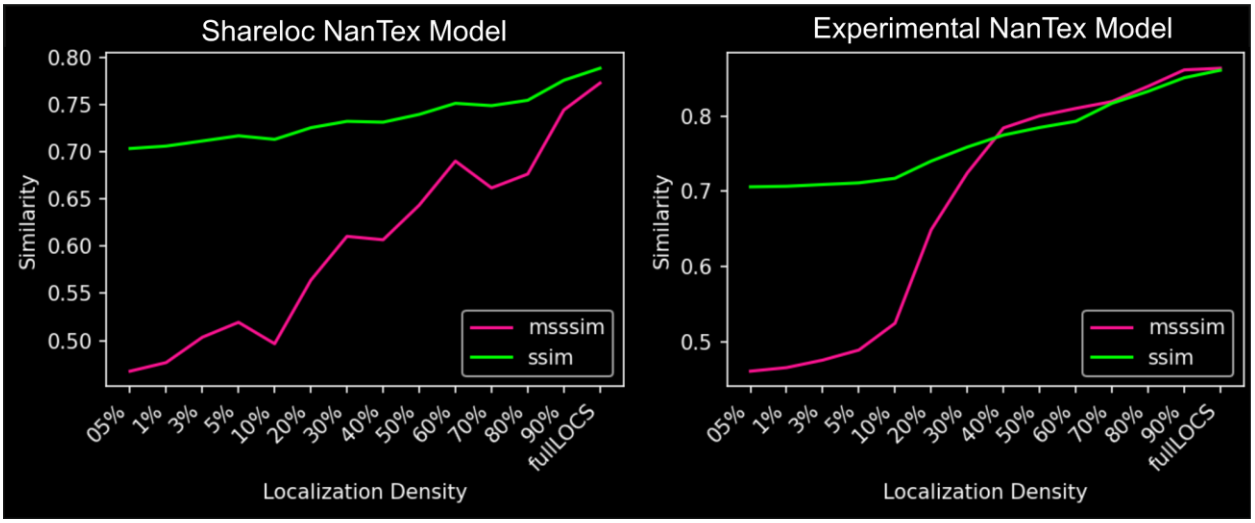


**Supplementary Figure 5 | Validation of *NanTex* recognition specificity using reduced localization density.** To test whether *NanTex* relies on true nanotextural features rather than global intensity or metadata, dSTORM microtubule datasets were systematically downsampled to simulate decreasing localization densities (x-axis). Reconstructions were evaluated against the corresponding full-density ground truth using SSIM (green) and MS-SSIM (magenta). Both the Shareloc-trained model (left) and the experimentally trained *NanTex* model (right) showed progressive breakdown of fidelity with decreasing localization density, particularly evident in MS-SSIM, which is more sensitive to fine-scale texture loss. At very low densities (≤10%), nanotexture features were no longer recognizable, confirming that *NanTex* recognition depends on genuine nanoscale structure. In contrast, reconstructions at ≥70% density retained high similarity to ground truth, demonstrating that sufficient localization density is essential for reliable demixing.

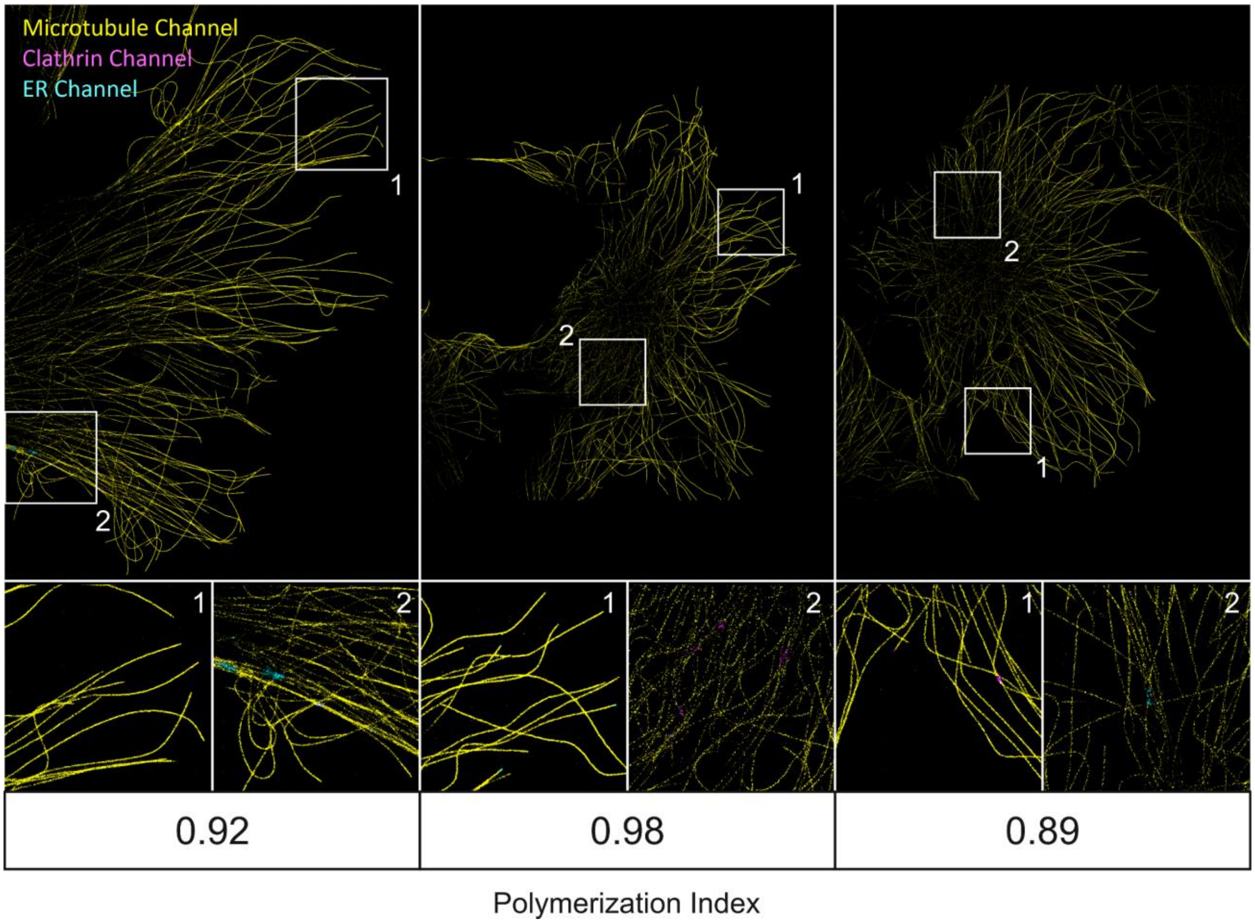


**Supplementary Figure 6 | High-purity microtubule *d*STORM datasets used as reference for quality control.** Three representative *d*STORM images of microtubules with near-maximal polymerization indices (PolIndex = 0.92, 0.98, and 0.89) are shown as gold-standard reference datasets for *NanTex* validation. These samples display high labeling efficiency and virtually no diffuse background, serving as benchmarks against which experimental variability and labeling quality can be assessed. *NanTex* demixed channels are color-coded (yellow: microtubules, cyan and magenta: other channels). Insets (1–2) highlight rare instances of apparent misallocation, underscoring the overall fidelity of the model in high-quality datasets. Together, these examples demonstrate that *NanTex* can act as a quantitative quality-control tool by comparing experimental preparations against benchmarked gold standards.

### Supplementary Videos

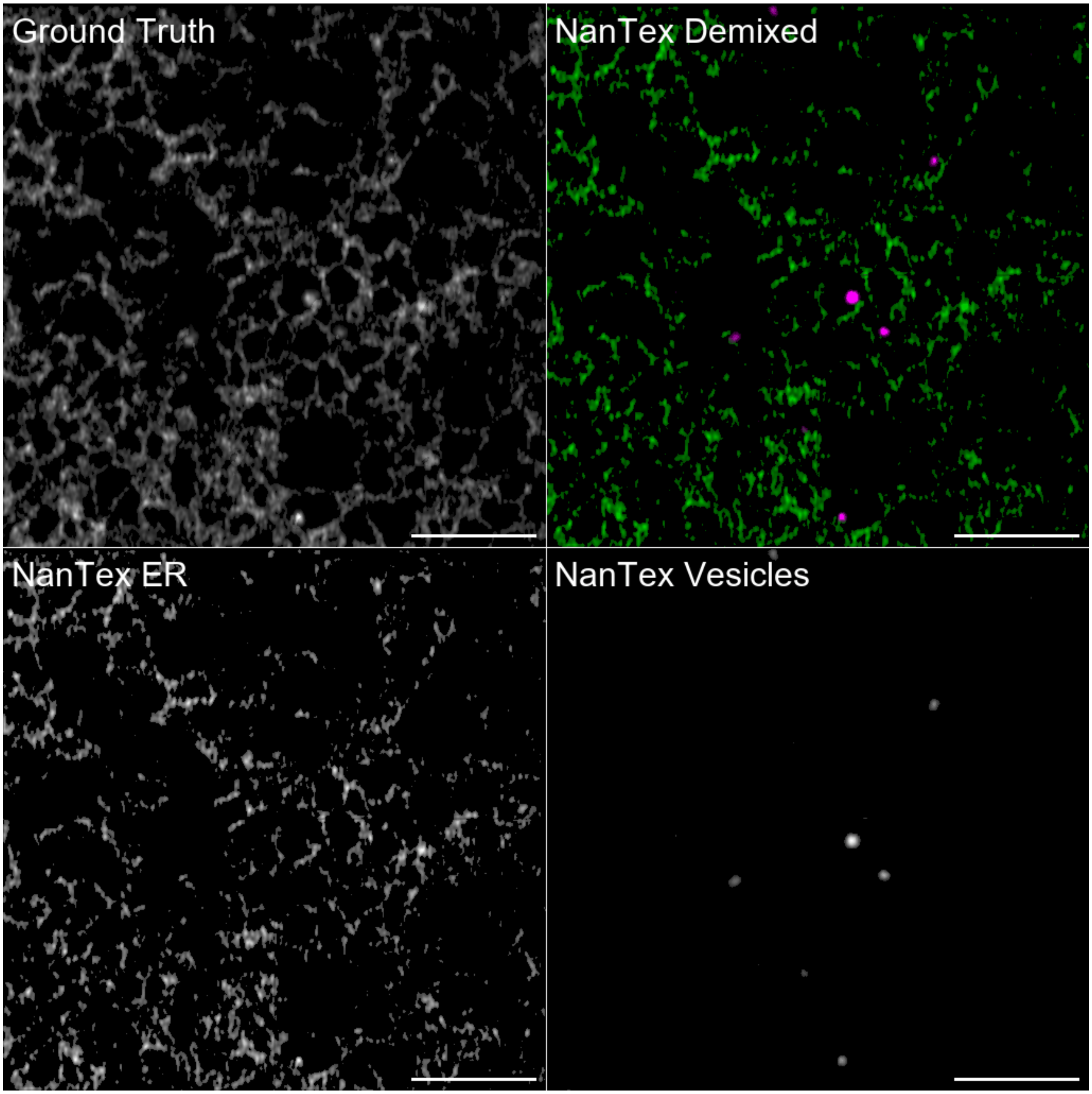


**Supplementary Video 1 | Live-cell Airyscan: ER–lysosome dynamics demixed by *NanTex*.** Time-resolved animated 2×2 panels illustrating *NanTex* ER-lysosomal multiplexing of Airyscan live-cell imaging. Upper left: original grayscale reference (ground truth). Upper right: *NanTex* demixed composite (ER = green, vesicles = magenta). Lower left: *NanTex* ER channel (feature 0, grayscale). Lower right: *NanTex* lysosome channel (feature 1, grayscale). This video highlights the ability of *NanTex* faithfully demixing ER-lysosomal interaction in live-cell imaging. Scale bars 5 µm.

Available as GIF.

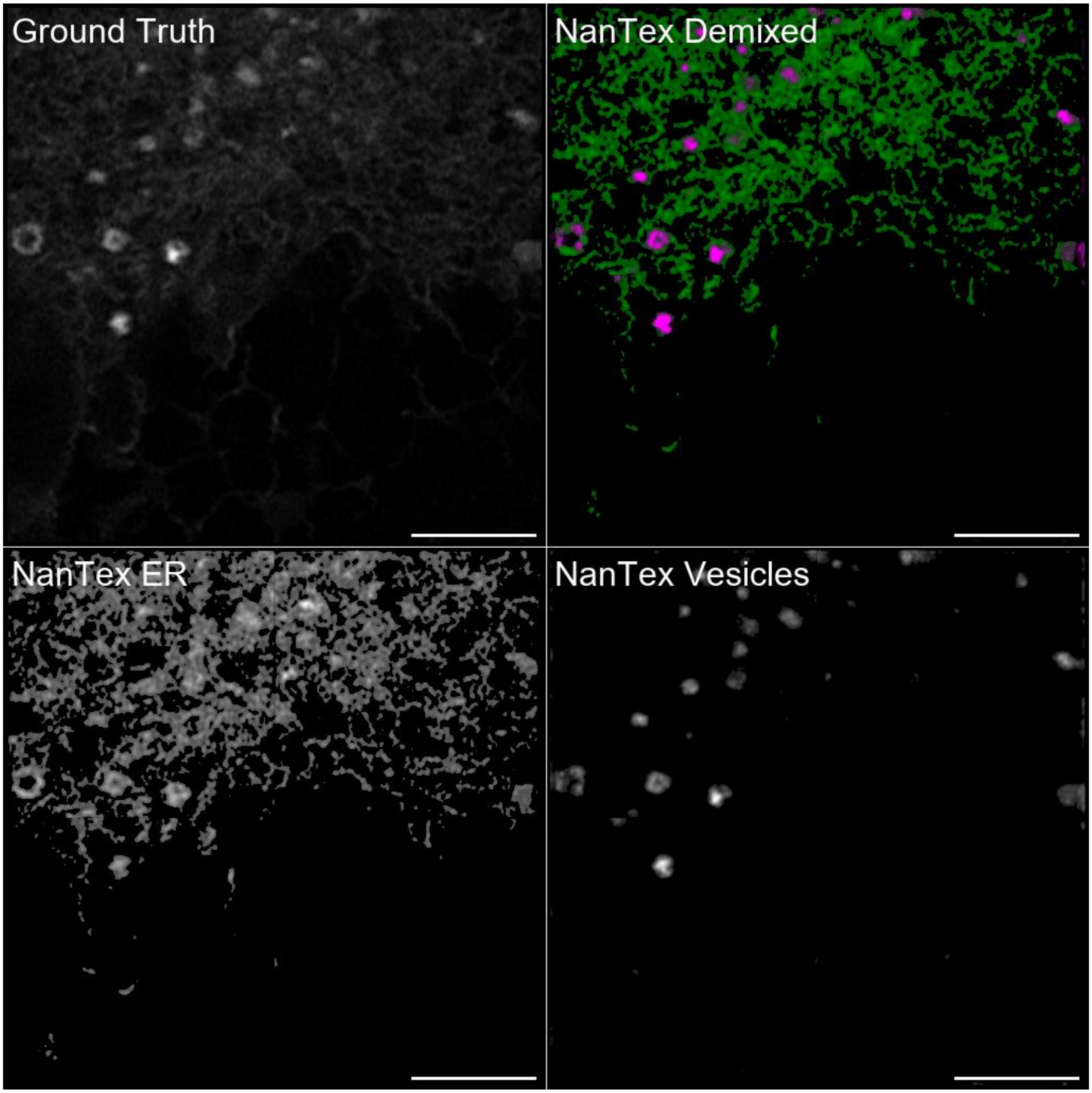


**Supplementary Video 2 | *NanTex* demixing of ER-only super-resolution Airyscan live-cell data.** Time-resolved animated 2×2 panels illustrating *NanTex* ER multiplexing of Airyscan live-cell imaging. Upper left: original grayscale reference (ground truth), contrast / dynamic range enhanced for visibility. Upper right: *NanTex* demixed composite (ER = green, vesicles = magenta). Lower left: *NanTex* ER channel (feature 0, grayscale). Lower right: *NanTex* vesicle channel (feature 1, grayscale). This video highlights the ability of *NanTex* demixing to resolve ER-associated structures and distinguish vesicular signals in live-cell multiplex experiments. Single particle tracking and MSD analysis is demonstrated in **Supplementary Fig. 4**. Scale bars 5 µm.

Available as GIF.
