## Supplementary figures and images for "NanTex enables computational multiplexing and phenotyping of organelles across super-resolution modalities"

### Supplementary Video 1

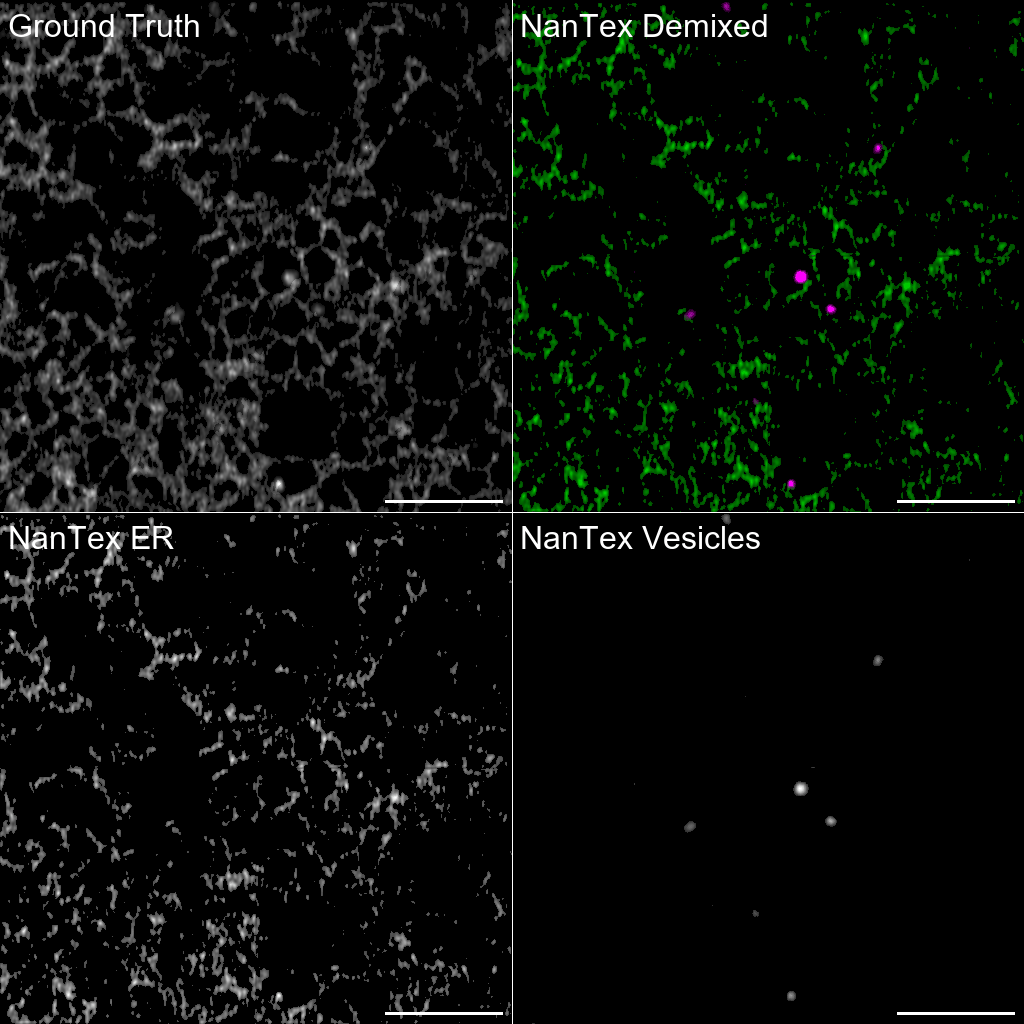

### Supplementary Video 2

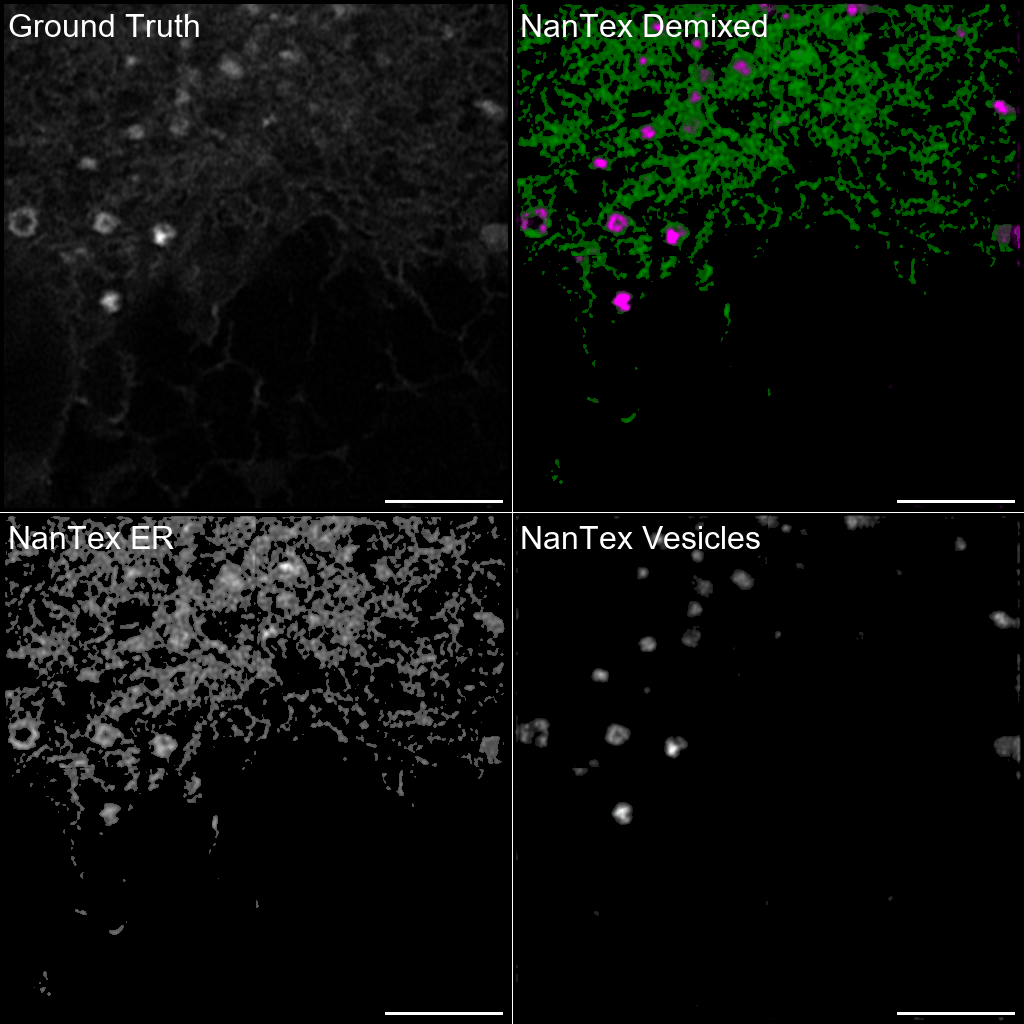
